# Promyelocytic Leukemia (PML) Nuclear Bodies (NBs) Induce Latent/Quiescent HSV-1 Genomes Chromatinization Through a PML-NB/Histone H3.3/H3.3 Chaperone Axis

**DOI:** 10.1101/217026

**Authors:** Camille Cohen, Armelle Corpet, Mohamed Ali Maroui, Olivier Binda, Nolwenn Poccardi, Antoine Rousseau, Pascale Texier, Nancy Sawtell, Marc Labetoulle, Patrick Lomonte

## Abstract

Herpes simplex virus 1 (HSV-1) latency establishment is tightly controlled by promyelocytic leukemia (PML) nuclear bodies (NBs) (or ND10), although their exact implication is still elusive. A hallmark of HSV-1 latency is the interaction between latent viral genomes and PML-NBs, leading to the formation of viral DNA-containing PML-NBs (vDCP-NBs). Using a replication-defective HSV-1-infected human primary fibroblast model reproducing the formation of vDCP-NBs, combined with an immuno-FISH approach developed to detect latent/quiescent HSV-1, we show that vDCP-NBs contain both histone H3.3 and its chaperone complexes, i.e., DAXX/ATRX and HIRA complex (HIRA, UBN1, CABIN1, and ASF1a). HIRA also co-localizes with vDCP-NBs present in trigeminal ganglia (TG) neurons from HSV-1-infected wild type mice. ChIP-qPCR performed on fibroblasts stably expressing tagged H3.3 (e-H3.3) or H3.1 (e-H3.1) show that latent/quiescent viral genomes are chromatinized almost exclusively with e-H3.3, consistent with an interaction of the H3.3 chaperones with multiple viral loci. Depletion by shRNA of single proteins from the H3.3 chaperone complexes only mildly affects H3.3 deposition on the latent viral genome, suggesting a compensation mechanism. In contrast, depletion (by shRNA) or absence of PML (in mouse embryonic fibroblast (MEF) *pml*^−/-^ cells) significantly impacts the chromatinization of the latent/quiescent viral genomes with H3.3 without any overall replacement with H3.1. Consequently, the study demonstrates a specific epigenetic regulation of latent/quiescent HSV-1 through an H3.3-dependent HSV-1 chromatinization involving the two H3.3 chaperones DAXX/ATRX and HIRA complexes. Additionally, the study reveals that PML-NBs are major actors in latent/quiescent HSV-1 H3.3 chromatinization through a PML-NB/histone H3.3/H3.3 chaperone axis.

**Author summary:** An understanding of the molecular mechanisms contributing to the persistence of a virus in its host is essential to be able to control viral reactivation and its associated diseases. Herpes simplex virus 1 (HSV-1) is a human pathogen that remains latent in the PNS and CNS of the infected host. However, the latency is unstable, and frequent reactivations of the virus are responsible for PNS and CNS pathologies. It is thus crucial to understand the physiological, immunological and molecular levels of interplay between latent HSV-1 and the host. Promyelocytic leukemia (PML) nuclear bodies (NBs) play a major role in controlling viral infections by preventing the onset of lytic infection. In previous studies, we showed a major role of PML-NBs in favoring the establishment of a latent state for HSV-1. A hallmark of HSV-1 latency establishment is the formation of PML-NBs containing the viral genome, which we called “viral DNA-containing PML-NBs” (vDCP-NBs). The genome entrapped in the vDCP-NBs is transcriptionally silenced. This naturally occurring latent/quiescent state could, however, be transcriptionally reactivated. Therefore, understanding the role of PML-NBs in controlling the establishment of HSV-1 latency and its reactivation is essential to design new therapeutic approaches based on the prevention of viral reactivation.

---

Herpes simplex virus 1 (HSV-1) is a human pathogen with neurotropic tropism and the causal agent of cold sores and more severe CNS pathologies such as encephalitis [1]. After the initial infection, HSV-1 remains latent in neuronal ganglia with the main site of latency being the trigeminal (or Gasserian) ganglion (TG). Two transcriptional programs are associated with HSV-1 infection, the lytic cycle and latency, which differ by the number and degree of viral gene transcription. The lytic cycle results from the sequential transcription of all viral genes (approximately 80) and leads to the production of viral progeny. The latency phase, occurring exclusively in neurons, is limited to the abundant expression of the so-called Latency Associated Transcripts (LATs), although physiologically a transitory expression of a limited number of lytic genes is not excluded, making latency a dynamic process [2–4].

Following lytic infection of epithelial cells at the periphery, the viral particle enters the axon termini of the innervating neurons by fusion of its envelope with the plasma membrane. The nucleocapsid is then carried into the neuron body by retrograde transport, most likely through the interaction of viral capsid components [5] with microtubule-associated proteins such as dynein and dynactin [6–10]. Once the nucleocapsid reaches the cell body, the virus phenotype changes from the one at the axon termini because most of the outer tegument proteins, including VP16, a viral transactivator that is essential for the onset of lytic infection, remain at the axonal tip [11–13]. Hence, when the viral DNA is injected into the neuron nucleus, it does not automatically benefit from the presence of VP16 to initiate lytic gene transcription. Rather, the balance between lytic and latent transcriptional programs most likely depends on stochastic events and on undescribed neuron-associated factor(s) able to initiate the transcription of VP16 through the activation of neuro-specific sequences present in the VP16 promoter [14]. Without VP16 synthesis, transcription of the viral genes encoding ICP4, the major transactivator protein, and ICP0, a positive regulator of viral and cellular gene transcription, is hampered. Hence, ICP4 and ICP0 gene transcription is unlikely to reach the required level to produce these two proteins above a threshold that would favor onset of the lytic cycle. Therefore, in neurons, commitment of the infectious process towards the lytic cycle or latency depends on a race between opposing infection-prone viral components and cellular features with antiviral activities.

Promyelocytic leukemia (PML) nuclear bodies (NBs) (also called ND10) are proteinaceous entities aimed to control viral infection as part of the cell and nucleus-associated intrinsic antiviral response but also through innate immunity associated with the interferon (IFN) response. Our recent studies have shown that PML-NBs tightly associate with incoming HSV-1 genomes in the nucleus of infected TG neurons in mouse models and in primary TG neuron cultures [15,16]. Hence, PML-NBs reorganize in structures called viral DNA-containing PML-NBs (vDCP-NBs), which are formed at early times during the process of HSV-1 latency establishment and persist during latency *per se* in a large subset of latently infected neurons in a mouse model of infection [15]. HSV-1 genomes trapped in the vDCP-NBs are transcriptionally repressed for LAT production [15]. It is known that HSV-1 latency, at least in the mouse model and possibly in humans, is heterogeneous at the single neuron level for the expression of LAT [15,17–24]. Therefore, although at the entire TG level HSV-1 latency could be a dynamic process from a transcriptional perspective, at the single neuron level, a strict, transcriptionally silent, quiescence can be observed, and vDCP-NB-containing neurons are major contributors of this latent/quiescent HSV-1 state. In humans, vDCP-NB-like structures have also been observed in latently infected TG neurons [16], suggesting that vDCP-NBs are probably molecular hallmarks of the HSV-1 latency process, including in the natural host.

Another essential feature of HSV-1 latency is the potent chromatinization of its 150-kb genome, which enters the nucleus of the infected cells as a naked/non-nucleosomal dsDNA [25–27]. Once the viral genome is injected into the nucleus of the infected neuron, it circularizes, associates with nucleosomes to become chromatinized, and remains as an episome that is unintegrated into the host cell genome [28]. Although latent viral genomes sustain epigenetic regulation, essentially through post-translational modifications of associated histones {Kubat:2004ty, Wang:2005vo, Knipe:2008uo, [29,30], not much is known about the mechanisms that induce their chromatinization and which specific histone variants are associated with these latent genomes. In mammals, specific H3 histone variants that differ by a few amino acids can influence chromatin compaction and transcriptional activity of the genome. The histone variant H3.3, a specific variant of the histone H3 that is expressed throughout the cell cycle, is deposited in a replication-independent manner, in contrast to H3.1 [31] (and for review [32]). Interestingly, death domain associated protein 6 (DAXX) and α-thalassemia mental retardation X-linked protein (ATRX), initially identified as a transcriptional repressor and a chromatin remodeler, respectively, are constitutively present in PML-NBs and have now been identified as H3.3-specific histone chaperones. The other histone H3.3 specific chaperone complex is called the HIRA complex, which is composed of Histone cell cycle regulator (HIRA), Ubinuclein 1 (UBN1), Calcineurin-binding protein 1 (CABIN1), and Anti-silencing function protein 1 homolog A (ASF1a) [31]. The HIRA complex does not normally accumulate in PML-NBs except upon entry of the cell into senescence [33,34]. The histone variant H3.3 itself localizes in PML-NBs in proliferating and senescent cells, linking PML-NBs with the chromatin assembly pathway independently of replication [35–37]. Because vDCP-NBs contain DAXX and ATRX [15,16,38], their involvement in the chromatinization of incoming HSV-1 genomes and/or long-term maintenance of chromatinized HSV-1 genomes is thus plausible.

Human primary fibroblasts or adult mouse primary TG neuron cultures infected through their cell body with a replication-defective HSV-1 virus, *in*1374, which is unable to synthesize functional ICP4 and ICP0 under specific temperature conditions, enable the establishment of a latent/quiescent state for HSV-1 [16,38–40]. The latent/quiescent state of HSV-1 in human primary fibroblasts has also been reproduced using engineered HSV-1 unable to express major immediate early genes [41,42]. We have shown that this latent/quiescent state is linked to the formation of vDCP-NBs, mimicking, at least concerning this particular structural aspect, the latency observed in a subset of neurons in mouse models and in humans [15,16]. Here, using the *in*1374-based *in cellula* model of infection, we showed that vDCP-NBs contained not only the DAXX and ATRX proteins but also all the components of the HIRA complex and H3.3 itself. HIRA was also detected co-localizing with vDCP-NBs in neurons of TG harvested from HSV-1 wild type infected mice. Both DAXX/ATRX and HIRA complex components were found interacting with multiple viral loci by chromatin immunoprecipitation (ChIP). Using the same approaches, we showed that latent/quiescent viral genomes were almost exclusively chromatinized with H3.3. Most interestingly, we found that H3.3 chromatinization of the viral genomes was dependent on intact PML-NBs, demonstrating that PML-NBs contribute to an essential part of the chromatinization of the latent/quiescent HSV-1 genomes. Overall, this study shows that the chromatinization of latent HSV-1 involves a PML-NB/histone H3.3/histone H3.3 chaperone axis that confers and probably maintains epigenetic marks on viral genomes.

## Results

### The HIRA complex components accumulate in the vDCP-NBs

The formation of vDCP-NBs is a molecular hallmark of HSV-1 latency, and vDCP-NBs are present in infected neurons from the initial stages of latency establishment to latency *per se* in mouse models [15,16]. Using a previously established *in vitro* latency system [39] consisting of human primary fibroblast cultures infected by a replication-deficient virus (hereafter called *in1374*) unable to express functional VP16, ICP4 and ICP0, we and others were able to reproduce the formation of vDCP-NBs [16,38]. We first verified that vDCP-NBs induced in human foreskin fibroblast (BJ) and other human primary cells infected by *in*1374 at a non-permissive temperature of 38.5°C, contained, in addition to PML, the proteins constitutively found in the PML-NBs, i.e., Sp100, DAXX, ATRX, SUMO-1 and SUMO-2/3 (Fig. S1Ai to vi, and Table S1). The DAXX/ATRX complex is one of the two chaperones of the histone variant H3.3 involved in the replication-independent chromatinization of specific, mostly heterochromatic, genome loci [43]. Interestingly, HSV-1 enters the nucleus of the infected cell as a naked/non-nucleosomal dsDNA and remains during latency as a circular chromatinized episome unintegrated in the host genome [28,44]. It is thus tempting to speculate that the presence of DAXX/ATRX in the vDCP-NBs could be linked to a process of initiation and/or maintenance of chromatinization of the latent/quiescent viral genome. The other H3.3 chaperone is known as the HIRA complex and was initially described as specific for the replication-independent chromatinization of euchromatin regions [31,45]. Remarkably, proteins of the HIRA complex are able to bind in a sequence-independent manner to a naked/non-nucleosomal DNA [46], suggesting that the HIRA complex could also participate in the recognition and chromatinization of the incoming naked HSV-1 genome. We thus investigated the localization of all members of the HIRA complex and found that they co-localize with the latent/quiescent HSV-1 genomes at 2 days post-infection (dpi) in BJ and other human primary cells (Figure 1 Ai to iv, Table S1). To confirm that the co-localization of members of the HIRA complex with the latent/quiescent HSV-1 could be reproduced in neuronal cells, adult mouse TG neuron cultures were infected with *in1374* for 2 days before performing immuno-FISH. Mouse Hira, which was the only protein of the HIRA complex detectable in mouse cells, showed a clear co-localization with a subset of viral genomes (Fig. 1B). To analyze whether this co-localization was also reproducible *in vivo*, immuno-FISH was performed on TG samples from HSV-1-infected mice. Hira was found to co-localize with HSV-1 genomes with the “multiple acute”/vDCP-NB pattern (see [16,47] in TG neurons from infected mice at 6 dpi (Fig. 1C) but not with the “single”/vDCP-NB pattern (see [15,47] at 28 dpi (Fig. 1D), suggesting a dynamic association of this protein with the vDCP-NBs.

**Figure 1.**
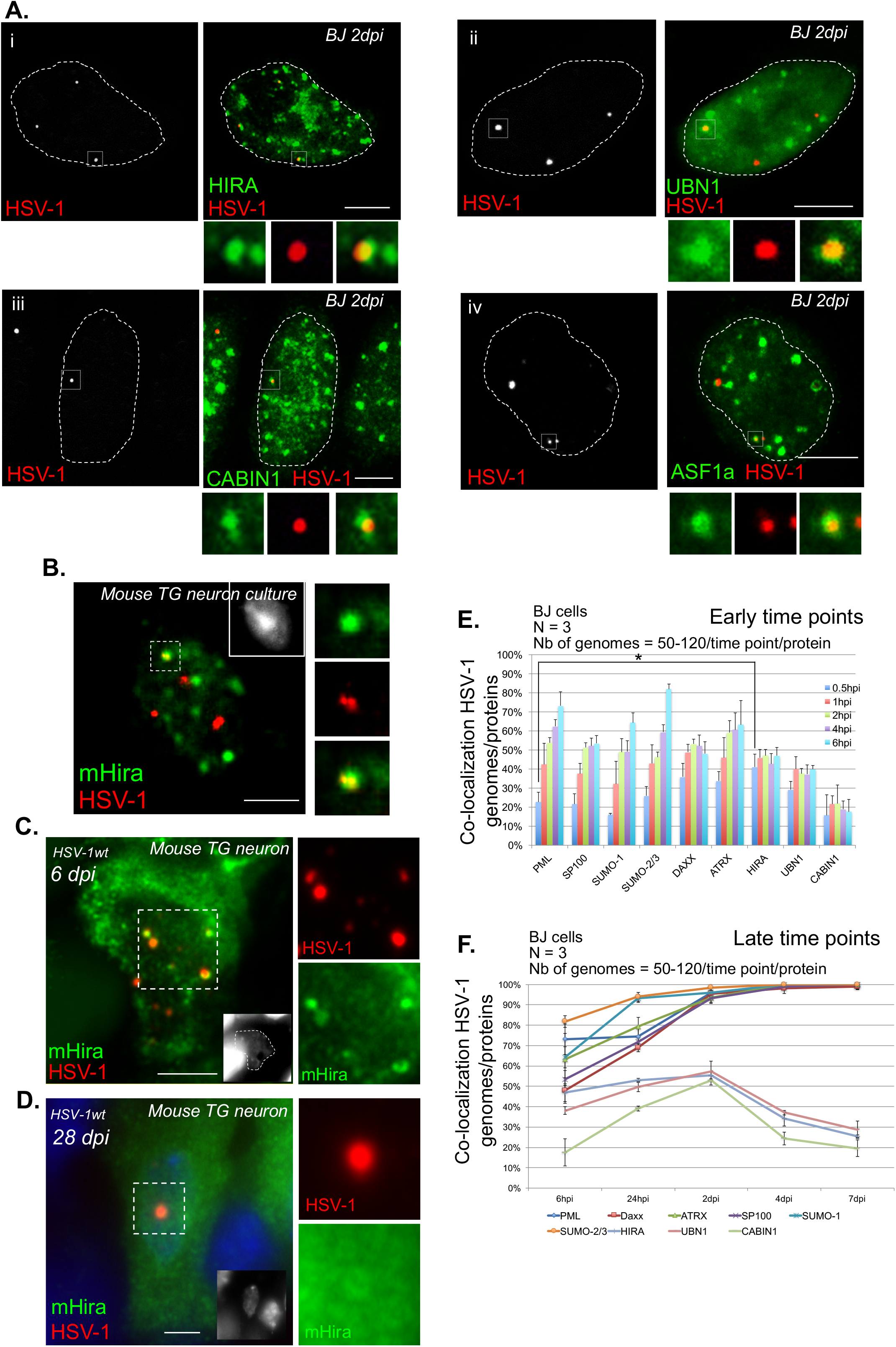
Latent/quiescent HSV-1 genomes co-localize with the HIRA complex. (A) Data from immuno-FISH experiments performed in human primary fibroblasts (BJ cells) infected for 2 days with the replication-defective HSV-1 virus *in*1374. HIRA (i), UBN1 (ii), CABIN1 (iii), ASF1a (iv) (green), and HSV-1 genomes (red) were detected. Scale bars = 5 µm. (B) Data from immuno-FISH experiments performed in adult mouse primary TG neuron cultures infected for 2 days with *in*1374 co-detecting mouse Hira (mHira, green) and HSV-1 genomes (red). Scale bars = 5 µm. (C) Data from immuno-FISH experiments performed in TG neurons of 6-day HSV-1wt-infected mice co-detecting mHira (green) and HSV-1 genomes (red). Nuclei (inset) were detected with DAPI (gray, blue). Scale bars = 10 µm. (D) Data from immuno-FISH experiments performed in TG neurons of 28-day HSV-1wt-infected mice co-detecting mHira (green) and HSV-1 genomes (red). Nuclei (inset) were detected with DAPI (gray, blue). Scale bars = 10 µm. (E) Quantifications from immuno-FISH experiments performed in infected BJ cells at early times post-infection with *in*1374 and representing the percentage of co-localization between incoming HSV-1 genomes and representative proteins of the PML-NBs (PML, Sp100, SUMO-1, SUMO 2/3) or H3.3 chaperone complex proteins (DAXX, ATRX, HIRA, UBN1, CABIN1). Data represent means from three independent experiments ± SD. The Student’s *t*-test was applied to assess the significance of the results. * = p< 0.05 (see Table S2 for data). (F) Quantifications from immuno-FISH experiments performed in infected BJ cells at late times post-infection with *in*1374 and representing the percentage of co-localization between incoming HSV-1 genomes and representative proteins of the PML-NBs (PML, Sp100, SUMO-1, SUMO 2/3) or H3.3 chaperone complex proteins (DAXX, ATRX, HIRA, UBN1, CABIN1). Data represent the means from three independent experiments ± SD (see Table S3 for data).

To analyze this dynamic association, co-localizations between incoming HSV-1 genomes and proteins of the PML-NBs or of the HIRA complex were quantified at early times from 30 min pi to 6 hpi using a synchronized infection procedure (Fig. 1E and Table S2). Except for the proteins of the HIRA complex, the percentages of co-localization increased with time. Interestingly, at 30 min pi, the percentage of co-localization of HSV-1 genomes with HIRA was significantly higher than with PML (41±7% vs 23±5%, p value = 0.03, Student’s *t-test*, Table S2). Although DAXX and ATRX also showed, on average, a greater percentage of co-localization with HSV-1 genomes (36±7% and 34±5% at 30 min, respectively) compared with PML, the data were not significant (Table S2). The absence of co-localization of mouse Hira with viral genomes with the “single”/vDCP-NB pattern in mouse TG neurons at 28 dpi suggests that longer infection times could lead to loss of proteins of the HIRA complex from the vDCP-NBs. Infection of BJ cells were reiterated as above, but this time quantifications were performed from 24 hpi to 7 dpi. Strikingly, whereas all the proteins permanently present in the PML-NBs remained co-localized with a maximum of 100% of the latent/quiescent HSV-1 genome from 2 dpi until 7 dpi, proteins of the HIRA complex peaked at 2 dpi, and then their co-localization decreased at longer times pi, confirming the temporary association of the HIRA complex with the vDCP-NBs (Figure 1F, and Table S3).

To definitively show that proteins of the HIRA complex were present in vDCP-NBs, immuno-FISH were performed on BJ cells infected for 2 days with *in1374* to detect either member of the HIRA complex, HSV-1 genomes, and PML. Strikingly, whereas in non-infected cells proteins of the HIRA complex showed predominant nucleoplasmic staining (Fig. 2i, iii, v, vii), in infected cells all the proteins clearly and systematically accumulated in PML-NBs (Fig. 2ii, iv, vi, viii). Consequently, HIRA, UBN1, CABIN1 and ASF1a co-localized with the latent/quiescent HSV-1 genomes in vDCP-NBs (arrows in Fig. 2ii, iv, vi, viii). Altogether, these data show that both DAXX/ATRX and HIRA complexes are present within vDCP-NBs in neuronal and non-neuronal cells, suggesting a role for these two complexes in latent/quiescent HSV-1 chromatinization.

**Figure 2.**
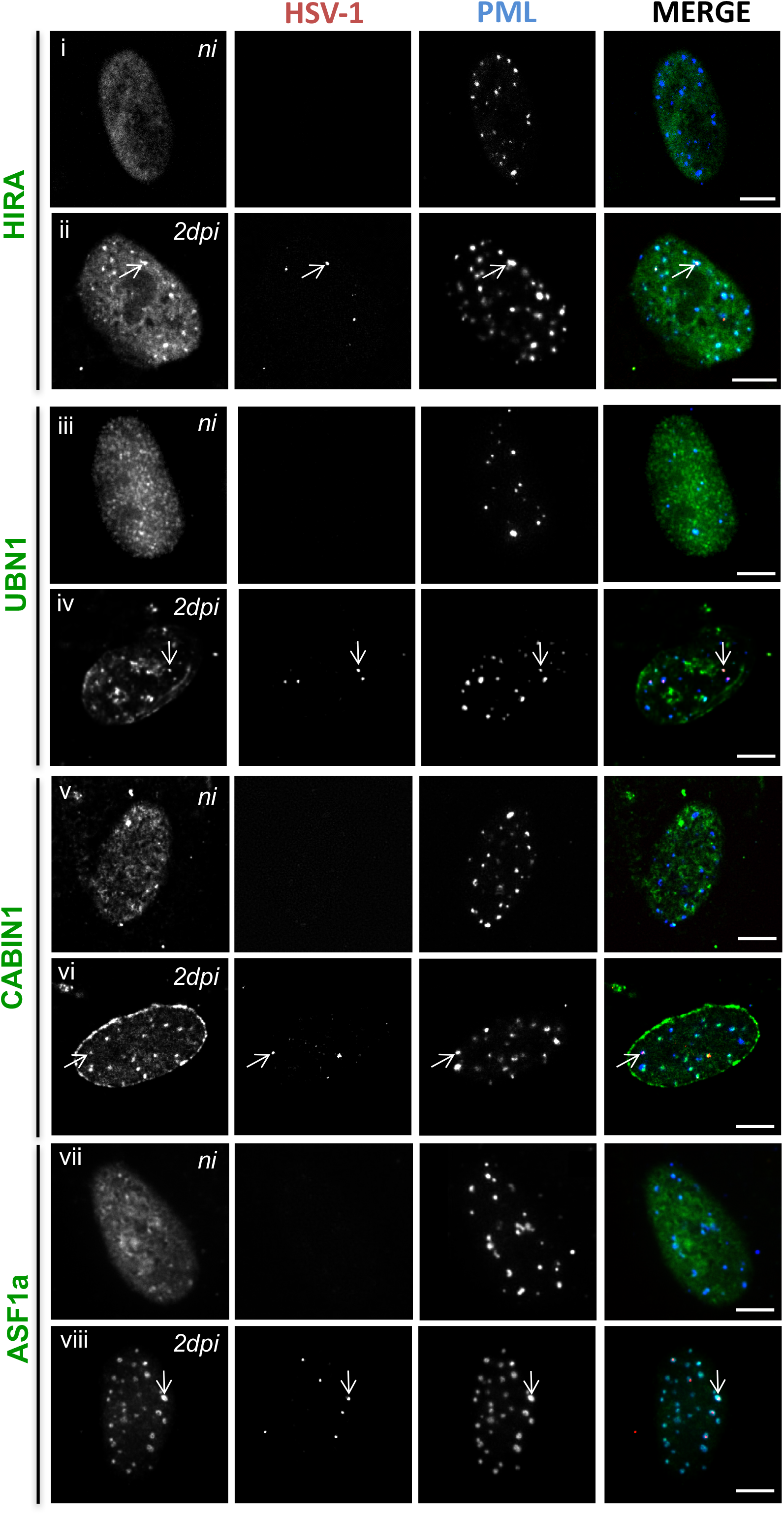
HSV-1 infection induces the accumulation of HIRA complex proteins in PML-NBs and co-localization with latent/quiescent HSV-1 genomes in vDCP-NBs. Immuno-FISH performed in BJ cells not infected (ni) (i, iii, v, vii) or infected for 2 days (ii, iv, vi, viii) with *in*1374. HIRA (i and ii), UBN1 (iii and vi), CABIN1 (v and vi), ASF1a (vii and viii) (gray, green), HSV-1 genomes (gray, red), and PML (gray, blue) were detected. Arrows indicate examples of the detection of HIRA complex proteins in vDCP-NBs. Scale bars = 5 µm.

### Histone H3.3 chaperones interact with incoming viral genome

The co-localization of proteins of the DAXX/ATRX and HIRA complexes with the incoming HSV-1 genomes and their presence in the vDCP-NBs suggested an interaction of these proteins with the viral genome, which we tested by chromatin immunoprecipitation (ChIP) assays. Since DAXX, HIRA, and UBN1 antibodies were not efficient in the ChIP experiments, we constructed cell lines stably expressing myc-DAXX, HIRA-HA, or HA-UBN1 by transduction of BJ cells with the respective lentivirus-expressing vectors (Fig. S2). Cells were infected with *in*1374 at 38.5°C and harvested to perform ChIP-qPCR on multiple loci spread over the HSV-1 genome (Fig. 3A). Significant enrichments compared to controls were detected for HIRA and UBN1 on several loci, confirming the interaction of these proteins with latent/quiescent HSV-1 genomes. Although ATRX bound to the viral genome, DAXX was not found significantly enriched, but we cannot exclude an alteration of the chromatin binding capacities of the tagged protein. Overall the ChIP-qPCR data confirmed the interaction of proteins of both the DAXX/ATRX and HIRA complexes with incoming vDCP-NBs-associated HSV-1 latent/quiescent genomes.

**Figure 3.**
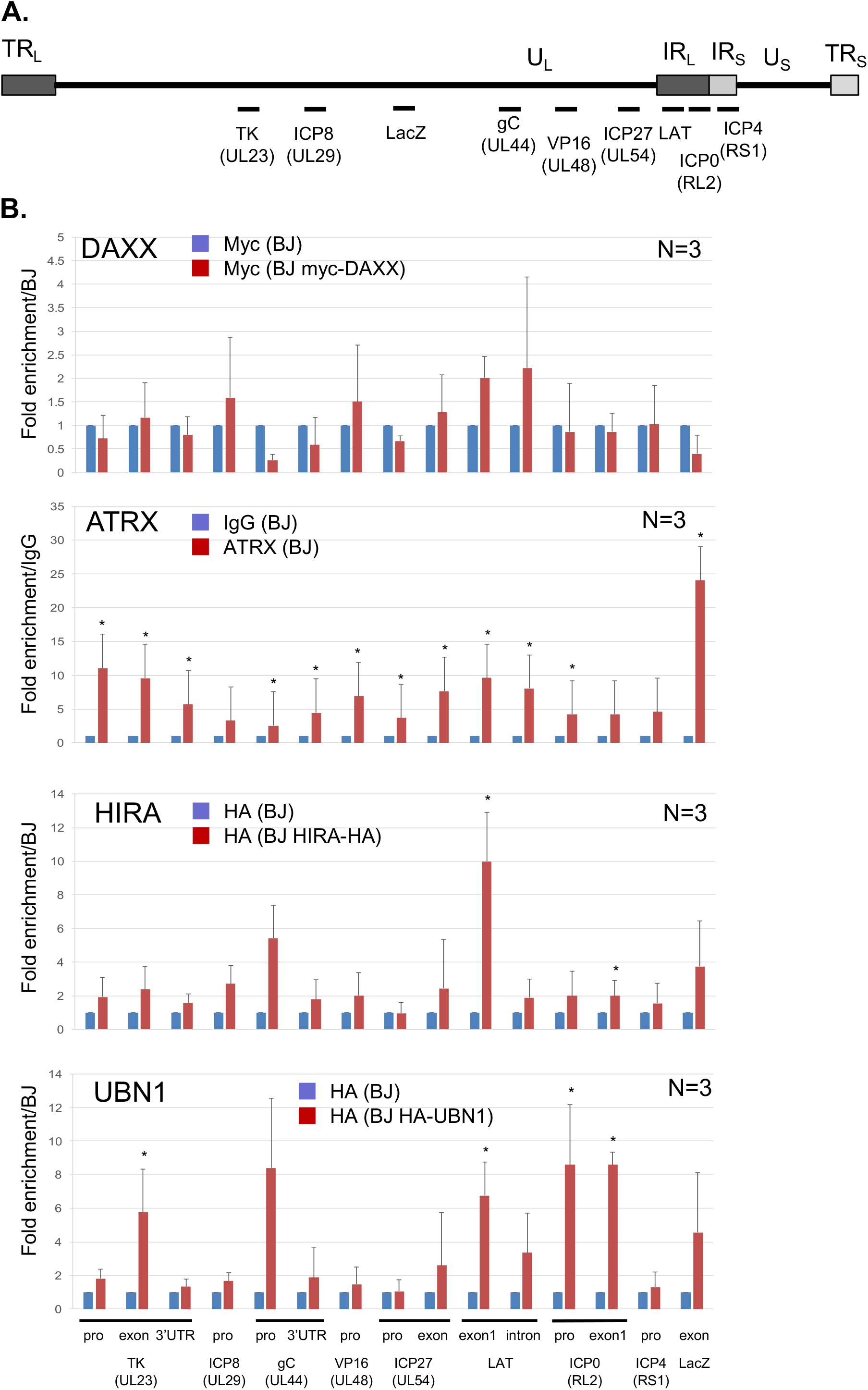
Components of the DAXX/ATRX and HIRA complexes associate with latent/quiescent HSV-1 genomes. (A) Schematic localization of the HSV-1 genome of the loci amplified by quantitative PCR. UL: Unit Long, US: Unit Short, TRL: Terminal Repeat Long, TRS: Terminal Repeat Short, IRL: Inverted Repeat Long, IRS: Inverted Repeat Short. (B) Chromatin immunoprecipitation (ChIP) associated with quantitative PCR (qPCR) performed in *in*1374-infected normal BJ cells or *in*1374-infected BJ cells expressing tagged versions of DAXX, HIRA, or UBN1. Anti-myc (DAXX) or anti-HA (HIRA and UBN1) antibodies were used. For ATRX, a native antibody was used, and the results were compared to ChIP with IgG as control. Data represent means from three independent experiments ± SD. Student’s *t*-test was applied to assess the significance of the results. * = p< 0.05.

### H3.3 is present in the vDCP-NBs and interacts with latent/quiescent HSV-1 genomes

The co-localization of the two histone H3.3 chaperone complexes with viral genomes suggested the chromatinization of HSV-1 latent genome with the histone variant H3.3. Histones H3.1 and H3.3 differ by only 5 amino acids, and, in our hands, no suitable antibody is available that can distinguish both histones by IF or IF-FISH. We constructed lentivirus-transduced BJ cell lines expressing a tagged version of either histone (e-H3.1 and e-H3.3) (see Materials and Methods, and [36], Fig. S3A and B). We confirmed that ectopic expression of e-H3.3 led to its accumulation in PML-NBs unlike e-H3.1 (Fig. S3C) [35,36]. *In*1374 infection of BJ e-H3.1/3-expressing cells led to the co-localization of viral genomes almost exclusively with e-H3.3 (Fig. 4Ai and ii, and B). Importantly, e-H3.3 co-localized with HSV-1 genomes together with PML in vDCP-NBs (Fig. 4C). The lack of co-localization of viral genomes with e-H3.1 was in accordance with the absence of detection of either H3.1 CAF-1 chaperone subunits (p150, p60, p48) in the vDCP-NBs (Fig. 4D, Table S1). To confirm that e-H3.3, unlike e-H3.1, interacted with HSV-1 genomes, ChIP experiments followed by qPCR were conducted on the same loci than those analyzed above. As expected, e-H3.3, but not e-H3.1, was highly enriched on the viral genome independently of the examined locus (Fig. 4E). To confirm that this discrepancy between e-H3.3 and e-H3.1 binding to viral genomes was not due to the ectopic expression of either protein, we performed a similar experiment using antibodies against native proteins. One specific antibody for H3.3 and working in ChIP experiments has been described [48]. Thus, we performed ChIP using antibodies against native H3.1/2 or H3.3 in normal BJ, or BJ e-H3.3 cells infected for 24 h by *in*1374. The results were similar to those obtained in infected BJ e-H3.3 using the anti-HA antibody, with a slightly better efficiency of the anti-HA antibody (Fig. S4A and B). These data confirmed that no bias was introduced in the ChIP experiments due to the use of tagged histones, and that latent/quiescent HSV-1 genomes are chromatinized with H3.3.

**Figure 4.**
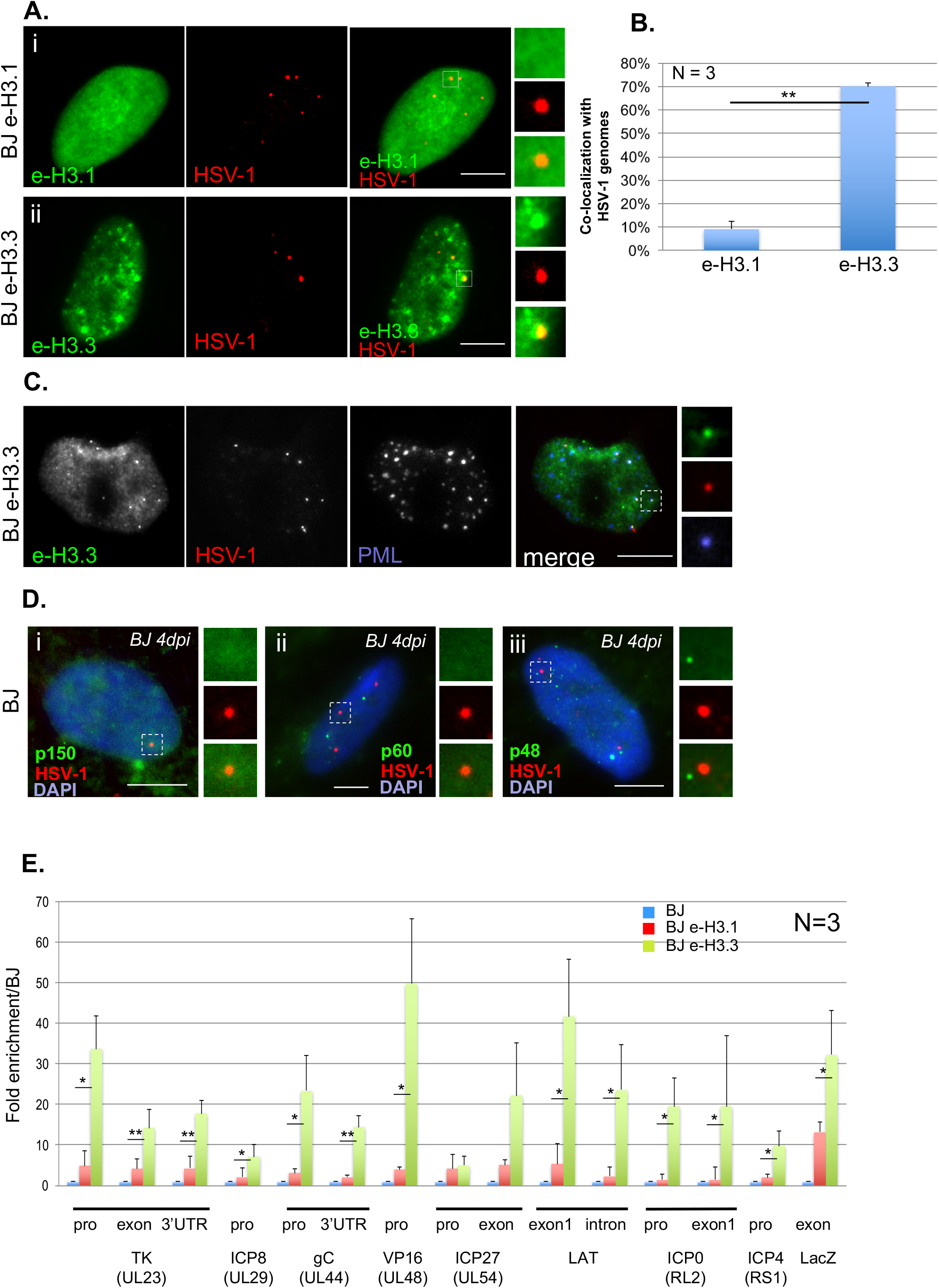
The histone variant H3.3 co-localizes and associates with latent/quiescent HSV-1 genomes. (A) Data from immuno-FISH experiments performed in e-H3.1 (i) or e-H3.3 (ii)-expressing BJ cells infected for 2 days with *in*1374. E-H3.1 or e-H3.3 (green), and HSV-1 genomes (red) were detected. Scale bars = 5 µm. (B) Quantification of the immuno-FISH experiments performed in (A). Data represent means from three independent experiments ± SD. The Student’s *t*-test was applied to assess the significance of the results. ** = p< 0.01. (C) Data from immuno-FISH experiments performed in e-H3.3-expressing BJ cells infected for 2 days with *in*1374. E-H3.3 (gray, green), HSV-1 (gray, red), and PML (gray, blue) were detected. Scale bars = 5 µm. (D) Data from immuno-FISH experiments performed in normal BJ cells infected for 2 days with *in*1374. H3.1 CAF chaperone complex proteins p150 (i), p60 (ii), and p48 (iii) (green), and HSV-1 genomes (red) were detected. Nuclei were detected with DAPI (blue). Scale bars = 5 µm. (E)ChIP-qPCR performed in *in*1374-infected normal BJ cells (blue) *in*1374-infected e-H3.1 (red) or e-H3.3 (green) expressing BJ cells. Anti-HA antibody was used for ChIP experiments. Analyzed viral loci were described previously. Data represent means from three independent experiments ± SD. The Student’s *t*-test was applied to assess the significance of the results. * = p< 0.05, ** = p< 0.01.

### Depletion of DAXX, ATRX, HIRA or UBN1 mildly affects HSV-1 genomes chromatinization with H3.3

To analyze the requirement of the histone H3.3 chaperones for the formation of the vDCP-NBs and HSV-1 chromatinization, DAXX, ATRX, HIRA or UBN1 were depleted by shRNAs in normal BJ cells or cells constitutively expressing e-H3.3 prior to infection with *in*1374 and completion of the experiments. The two tested shRNAs for each protein significantly diminished mRNA and protein quantities in BJ cells (Fig. S5A and B). None of the shRNA impacted the detection of PML-NBs, suggesting that PML-NBs were potentially functional in the absence of either protein (Fig. S6). We first measured the impact of the depletion of each protein on the co-localization of HSV-1 genomes with PML. Both shRNAs for each protein provided similar results, i.e., a significant decrease in the co-localization between HSV-1 genomes and PML and thus a decrease in the formation and/or stability of the vDCP-NBs (Fig. 5A and B, Table S4). The absence of HIRA had a much weaker effect compared to the three others. These data showed that the inactivation of either H3.3 chaperone complex by removal of one of its component affected to a certain extent the fate of vDCP-NBs suggesting a dependency between the activity of each H3.3 chaperone complex and the formation and/or maintenance of the vDCP-NBs.

**Figure 5.**
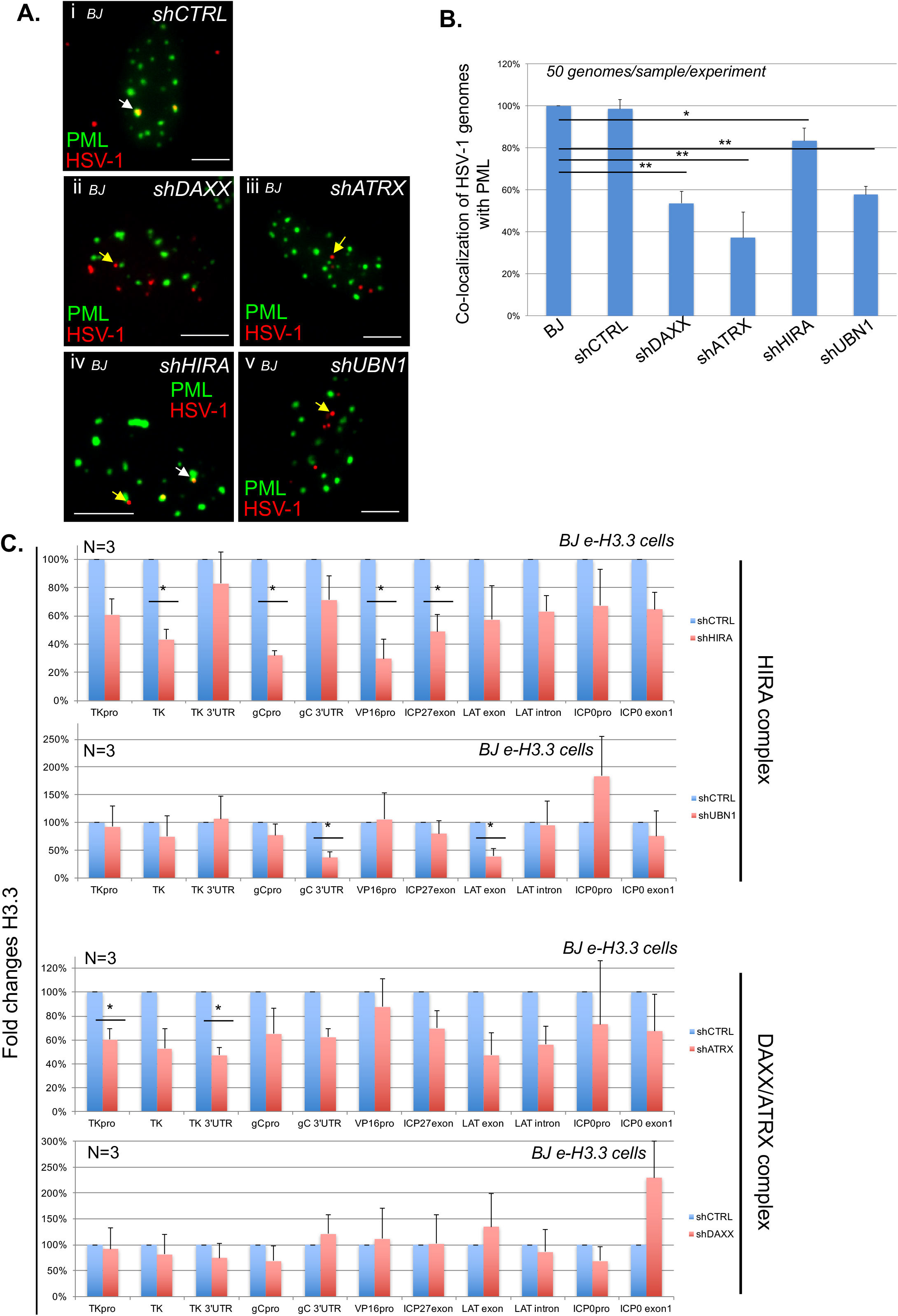
Depletion of DAXX, ATRX, HIRA, or UBN1 significantly affects the formation of vDCP-NBs, but only mildly affects the association of H3.3 with the latent/quiescent HSV-1 genome. Normal or e-H3.3-expressing BJ cells were first transduced with shRNA-expressing lentiviruses before analysis. (A) Data from immuno-FISH experiments performed in BJ cells infected with *in*1374 for 24 h. PML (green) and HSV-1 genomes (red) were detected in lentivirus-transduced BJ cells expressing control (shCTRL) or targeted shRNAs. Scale bars = 5 µm. (B) Quantifications of the immuno-FISH experiments performed in (A). Data represent means from three independent experiments ± SD. The Student’s *t*-test was applied to assess the significance of the results. * = p< 0.05, ** = p< 0.01 (see Table S4 for data). (C) ChIP-qPCR for the detection of e-H3.3 associated with HSV-1 genomes and performed in e-H3.3-expressing BJ cells infected with *in*1374 for 24 h and previously transduced with a lentivirus expressing a control shRNA (shCTRL, blue) or a targeted shRNA (red). Anti-HA antibody was used for the ChIP experiments. The analyzed viral loci were described previously. Data represent means from three independent experiments ± SD. The Student’s *t*-test was applied to assess the significance of the results. * = p< 0.05 (see Table S5 for data).

We then analyzed the potential impact of the loss of vDCP-NB stability on the H3.3-dependent HSV-1 chromatinization. We performed H3.3 ChIP in *in*1374-infected BJ e-H3.3 cells that had been previously depleted for DAXX, ATRX, HIRA or UBN1 using one of the previously validated shRNAs (Fig. S7A and B). ChIP-qPCR showed that the depletion of chaperones had overall only a mild impact on the number of loci showing a significant decrease in H3.3 association. Although depletion of HIRA had a more profound impact on H3.3 deposition than depletion of UBN1 or ATRX, the depletion of DAXX showed no significant effect (Fig. 5C, Table S5). Thus, the inactivation of individual components of each H3.3 chaperone complex, although having some impact on the association of some loci with H3.3, does not seem sufficient to completely abolish the process of HSV-1 chromatinization. These results suggest that ATRX/DAXX may compensate for the loss of HIRA and vice versa.

### PML-NBs are essential for H3.3 chromatinization of latent/quiescent HSV-1 genomes

The above experiments were conducted in a context where the cells, although deficient for the activity of one H3.3 chaperone complex at a time, still contained intact PML-NBs accumulating e-H3.3 (Fig. S6 and S8). Therefore, we hypothesized that the accumulation of H3.3 within the PML-NBs could be one of the key events acting upstream of the H3.3 chaperone complex activity for the induction of chromatinization of the latent/quiescent HSV-1 by H3.3. Thus, we analyzed the HSV-1 chromatinization in cells lacking PML-NBs. We had previously analyzed the impact of PML depletion on the co-localization of the DAXX/ATRX and HIRA chaperone complexes with the HSV-1 genomes. In a previous study conducted in HSV-1 latently infected PML KO mice, we showed that the absence of PML significantly impacted the number of latently infected TG neurons showing the “single”/vDCP-NB HSV-1 pattern and favored the detection of neurons containing the “multiplelatency” pattern prone to LAT expression [15,47]. We analyzed the very few neurons showing a “single”/vDCP-NB-like pattern in the latently infected PML KO mice for the co-localization of DAXX and ATRX with the viral genomes. We could not detect any of the two proteins co-localizing with the latent HSV-1 genomes (Fig. 6A i to vi). Although informative, these *in vivo* studies did not allow analysis of the real impact of the absence of PML on the co-localization of the other PML-NB-associated proteins with latent HSV-1 genomes because the neurons showing the “single”/vDCP-NB-like pattern were too few to quantify the effect. We thus depleted PML in normal BJ cells using a PML shRNA-expressing lentivirus transduction. We verified the efficiency of the shRNAs against PML in normal BJ cells by IF, RT-qPCR and WB (Figure S9A-C). PML-depleted BJ cells were superinfected with *in*1374, and immuno-FISH was performed at 2 dpi to analyze the co-localization of HSV-1 genomes with DAXX, ATRX, HIRA, and UBN1 (Fig. 6B). Both PML shRNAs provided similar results. The quantification of the data showed that, similarly to the *in vivo* situation, the depletion of PML significantly decreased the co-localization of DAXX and ATRX with latent/quiescent HSV-1 genomes, leaving HIRA and UBN1 unaffected for their co-localization (Fig. 6C, and Table S6). Thus, we analyzed whether the deficit of DAXX/ATRX co-localization with the latent/quiescent HSV-1 genomes as a consequence of the absence of PML-NBs could impact the chromatinization of HSV-1 with H3.3.

**Figure 6.**
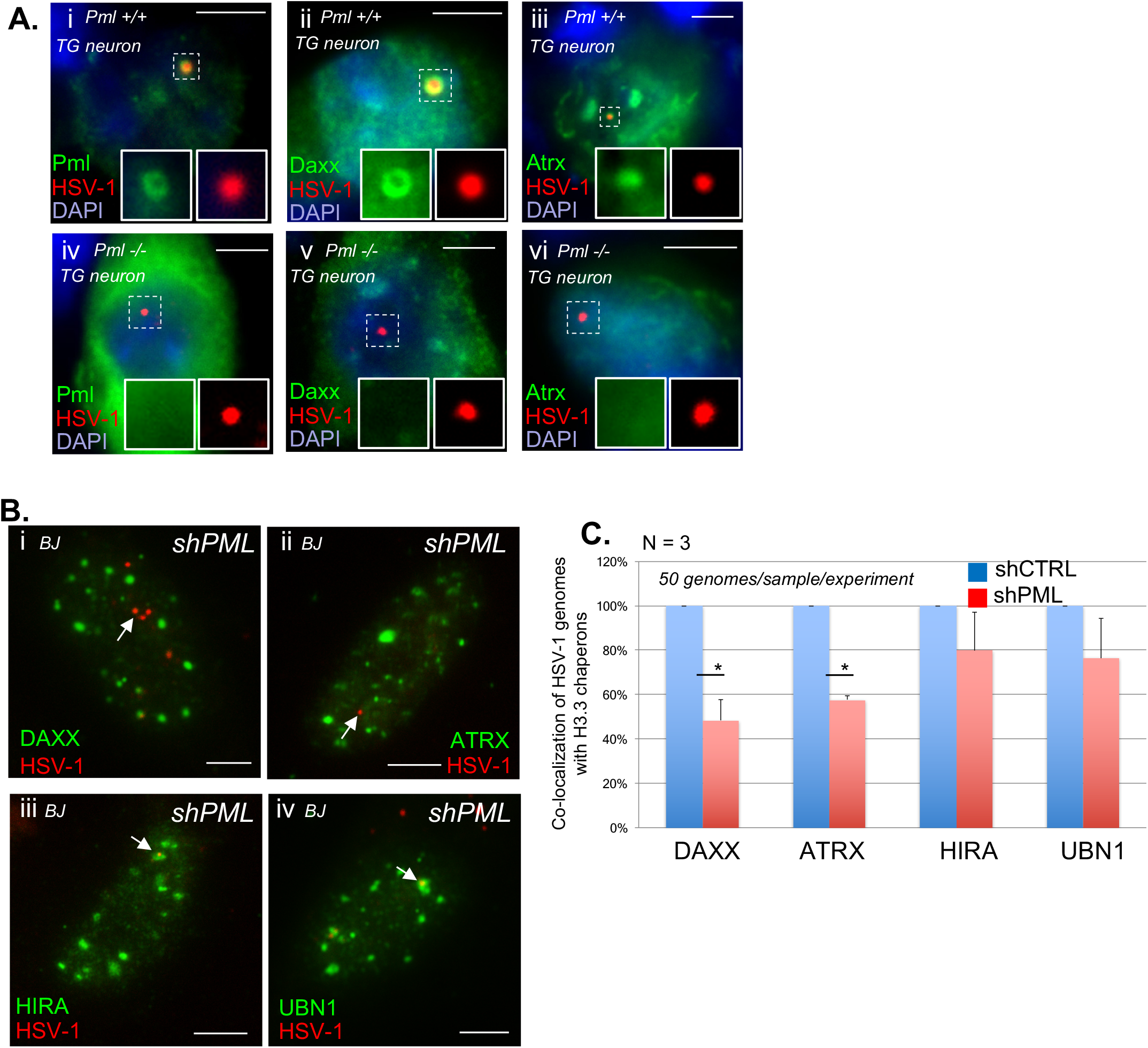
Absence of PML decreases the co-localization of DAXX and ATRX but not HIRA and UBN1 with latent/quiescent HSV-1 genomes. (A) Data from immuno-FISH experiments performed in TG tissues from *pml* ^+/+^ and *pml* ^−/-^ infected mice at 28 dpi. Pml, Daxx, Atrx (green), and HSV-1 genomes (red) were detected. Nuclei were detected with DAPI (blue). Scale bars = 10 µm. (B) Data from immuno-FISH experiments performed in BJ cells depleted of PML by transduction with a PML-targeted shRNA-expressing lentivirus and subsequently infected with *in*1374 for 2 days. DAXX, ATRX, HIRA or UBN1 (green), and HSV-1 genomes (red) were detected. Scale bars = 5 µ (C) Quantification of the immuno-FISH experiments performed in (B). Data represent means from three independent experiments ± SD. The Student’s *t*-test was applied to assess the significance of the results. * = p< 0.05 (See Table S7 for data).

We first generated BJ e-H3.3 cells depleted for PML by shRNA-expressing lentivirus transduction similarly to the BJ cells (Figure S9D-E). BJ e-H3.3 control or PML-depleted cells were superinfected with *in*1374 to perform immuno-FISH to analyze the co-localization of HSV-1 genomes with H3.3 (Fig. 7A). Quantification of the data clearly showed a significant decrease in the co-localization of latent/quiescent HSV-1 genomes with H3.3 compared with the controls (Fig. 7B), suggesting an impact of the absence of PML-NBs on the latent/quiescent HSV-1 association with H3.3. To complement these results at a more quantitative level, we performed ChIP-qPCR experiments on e-H3.3. The data clearly showed a significant impact of the absence of PML-NBs on the H3.3 association with multiple viral loci, with a depletion of H3.3 in all the analyzed loci excluding one (Fig. 7C, Table S7), which could not be due to an indirect effect of PML depletion on H3.3 stability because e-H3.3 protein levels were similar in control cells and cells depleted for PML (Fig. 7D). Both PML shRNAs provided similar results. To confirm that the absence of PML had an impact on the H3.3 association with latent/quiescent viral genomes, we performed ChIP on *in*1374-infected control mouse embryonic fibroblast (MEF) *pml*^+/+^ or MEF *pml*^−/-^ cells previously engineered by lentiviral transduction to express e-H3.3 (Fig. 7E). The data confirmed the deficit of an association of e-H3.3 with the latent/quiescent viral genomes in the absence of PML (Fig. 7F). Finally, we wanted to analyze whether the deficit of the H3.3 association with the viral genome in the absence of PML could be compensated by an increase of H3.1 on viral loci. The data from BJ e-H3.1 cells depleted for PML and infected with *in*1374 showed that H3.1 did not replace H3.3 on the same analyzed viral loci (Fig. S10). Altogether, these data demonstrate the essential role of PML-NBs, probably through the DAXX/ATRX complex activity, in the exclusive H3.3 chromatinization of incoming viral genomes forced to adopt a vDCP-NB-associated latent/quiescent pattern due to a deficit in the onset of lytic cycle.

**Figure 7.**
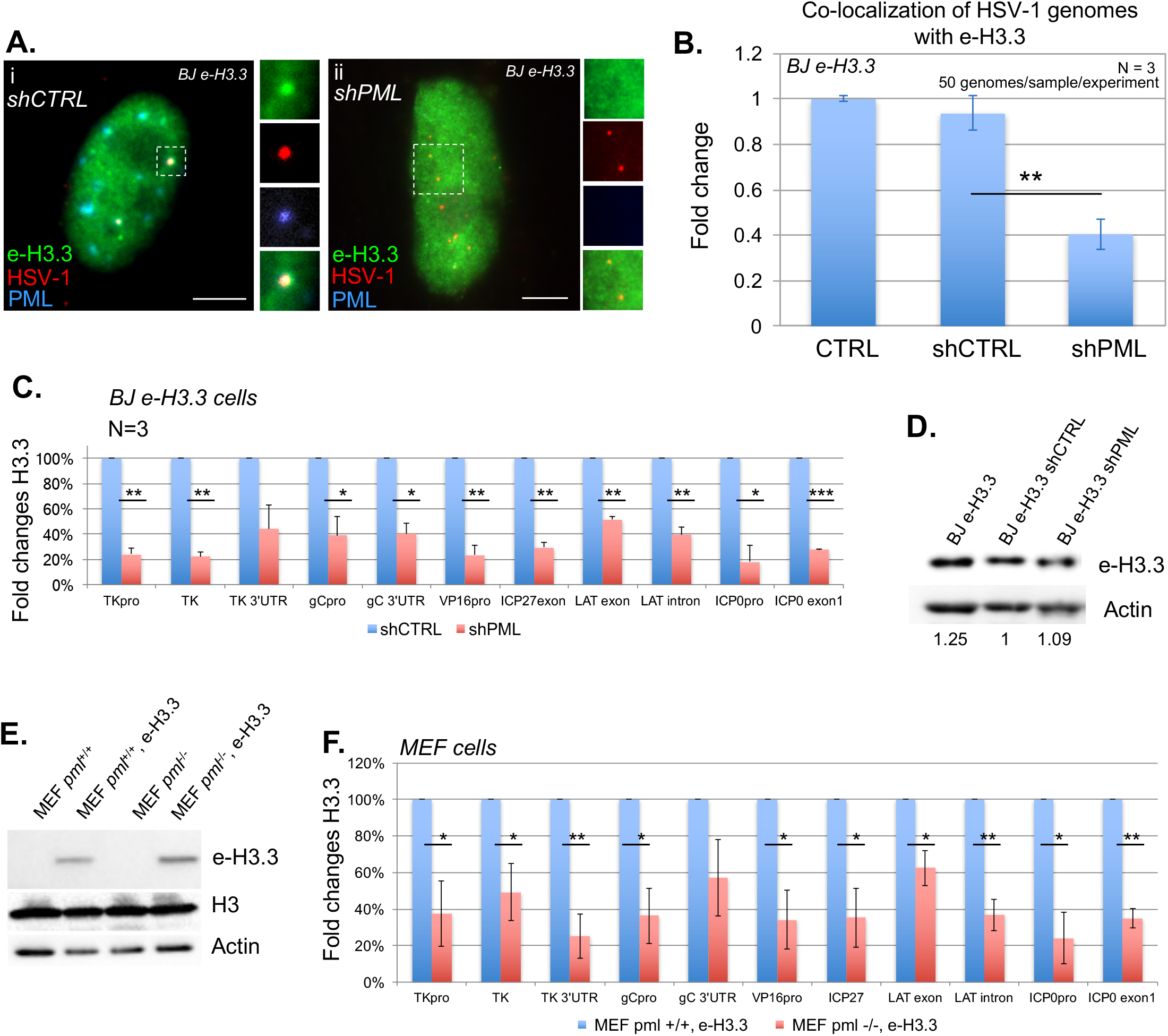
Depletion of PML significantly impacts the association of H3.3 with latent/quiescent HSV-1 genomes. (A) Data from immuno-FISH experiments performed in e-H3.3-expressing BJ cells transduced with a control (shCTRL) or PML (shPML) shRNA-expressing lentivirus and subsequently infected with *in*1374 for 2 days. E-H3.3 (green), HSV-1 genomes (red), and PML (blue) were detected. Scale bars = 5 µm. (B) Quantification of the immuno-FISH experiments performed in (A). Data represent means from three independent experiments ± SD. The Student’s *t*-test was applied to assess the significance of the results. ** = p< 0.01. (C) ChIP-qPCR for the detection of e-H3.3 associated with HSV-1 genomes and performed in e-H3.3-expressing BJ cells infected by *in*1374 for 24 h and previously transduced with a lentivirus expressing a control shRNA (shCTRL, blue) or a PML shRNA (shPML, red). Anti-HA antibody was used for the ChIP experiments. The analyzed viral loci were described previously. Data represent means from three independent experiments ± SD. The Student’s *t*-test was applied to assess the significance of the results. * = p< 0.05 (See Table S6 for data). (D) WB for the detection of e-H3.3 in control e-H3.3-expressing BJ cells (CTRL) or e-H3.3-expressing BJ cells transduced with a lentivirus expressing a control shRNA (shCTRL) or a PML shRNA (shPML). Actin was detected as a loading control. (E) WB for the detection of e-H3.3 in control and e-H3.3-expressing MEF *pml*^+/+^ or *pml* ^−/-^ cells. Actin and total histone H3 were detected as loading controls. (F) ChIP-qPCR for the detection of e-H3.3 associated with HSV-1 genomes and performed in e-H3.3-expressing *pml*^+/+^ (blue) or *pml*^−/-^ (red) MEF cells infected by *in*1374 for 24 h. Anti-HA antibody was used for the ChIP experiments. Analyzed viral loci were described previously. Data represent means from three independent experiments ± SD. The Student’s *t*-test was applied to assess the significance of the results. * = p< 0.05, ** = p< 0.01. See Table S8 for data.

## Discussion

The HSV-1 genome enters the nucleus of infected neurons, which support HSV-1 latency as a naked/non-nucleosomal DNA. Many studies have described the acquisition of chromatin marks on the viral genome concomitantly to the establishment, and during the whole process, of latency. Paradoxically, although it is undisputable that these chromatin marks will predominantly be associated with latency and reactivation, few data are available for the initiation of the chromatinization of the incoming viral genome. Here, we demonstrate the essential contribution of PML-NBs in the process of chromatinization of incoming HSV-1 genomes meant to remain in a latent/quiescent state. We showed that PML-NBs are essential for the association of the histone variant H3.3 with the latent/quiescent HSV-1.

Two members of the HIRA complex, HIRA and ASF1a, were previously shown to be involved in H3.3-dependent chromatinization of HSV-1 genomes at early times after infection in non-neuronal and non-primary cells favoring the onset of the lytic cycle [49,50]. Our *in vivo* data in TG neurons and *in vitro* data in infected human primary fibroblasts or adult mice TG neuron cultures, show that all the proteins of the HIRA complex accumulate within specific nucleoprotein structures called the viral DNA-containing PML-NBs or vDCP-NBs. vDCP-NB is a transcriptionally silent HSV-1 genome pattern that we previously demonstrated *in vivo* to be associated with the establishment of latency from the early steps of neuron infection [16]. Additionally, our data show that (i) the mouse Hira protein *in vivo*, and all the components of the HIRA complex in cultured cells, accumulate in vDCP-NBs temporarily, and (ii) significantly greater amount of incoming HSV-1 genomes co-localize with HIRA compared with PML at very early times pi (30 min). These data suggest that the HIRA complex could also be involved to some extent in the establishment of HSV-1 latency by the initial recognition of the incoming naked/non-nucleosomal viral DNA and the chromatinization of non-replicative HSV-1 genomes intended to become latent. In this respect, a recent study suggested an anti-viral activity associated with HIRA against HSV-1 and murine cytomegalovirus lytic cycles [51].

Interestingly, all the proteins of the HIRA complex have been previously shown to be able to directly bind to naked DNA in a sequence-independent manner, in contrast to DAXX and ATRX proteins [46]. Nevertheless, our ChIP data highlighted some specific viral genome loci that interact at least with ATRX. Thus, it is likely that ATRX, unlike HIRA and UBN1, indirectly binds to the viral genome. The gamma-interferon-inducible protein 16 (IFI16), a member of the PYHIN protein family, has been described as a nuclear sensor of incoming herpesviruses genomes and suggested to promote the addition of specific chromatin marks that contribute to viral genome silencing [52–59]. A proteomic study determining the functional interactome of human PYHIN proteins revealed the possible interaction between ATRX and IFI16 [60]. Thus, it will be interesting to determine in future studies if IFI16 and H3.3 chaperone complexes physically and functionally cooperate in the process of chromatinization of the latent/quiescent HSV-1 genome.

One of the main finding of our study is the demonstration of the implication of PML-NBs in the H3.3-dependent chromatinization of the latent/quiescent HSV-1 genomes. A close link between PML-NBs and H3.3 in chromatin dynamics has been demonstrated during oncogene-induced senescence (OIS). In OIS, expression of the oncogene H-RasV12 induces DAXX-dependent relocalization of neo-synthesized H3.3 in the PML-NBs before a drastic reorganization of the chromatin to form senescence-associated heterochromatin foci [35,36]. Hence, the implication of the PML-NBs in the deposition of H3.3 on specific cellular chromatin loci has also been reported [36,37]. The present study shows that the absence of Pml in HSV-1wt latently infected Pml KO mice, or the depletion of PML by shRNA in BJ cells infected with *in*1374, significantly affects the co-localization of DAXX and ATRX, but not HIRA and UBN1, with latent/quiescent HSV-1 genomes, confirming previous studies for DAXX/ATRX [38]. Taken together with the deficit of the H3.3 association with the viral genomes in the absence of PML-NBs, these data suggest that a significant part of the latent/quiescent HSV-1 genome chromatinization by H3.3 could occur through the activity of the DAXX/ATRX complex in association with the PML-NBs.

Given the particular structure formed by the latent/quiescent HSV-1 genome with the PML-NBs, our study raises the question of the possible acquisition of chromatin marks in the vDCP-NBs. The depletion of H3.3, which almost exclusively participates in latent/quiescent HSV-1 genome chromatinization, does not prevent the formation of vDCP-NBs and is rather in favor of such a scenario (Fig. S11). It is unlikely that a possible replacement of H3.3 by the canonical H3.1 for the chromatinization of the incoming HSV-1 genomes could occur prior to the association with vDCP-NBs. Indeed, our multiple immuno-FISH and ChIP assays failed to detect H3.1 and/or H3.1 chaperones that associate or co-localize with viral genomes. Nonetheless, we cannot rule out a possible replacement of H3.3 with another H3 variant.

Even if a process of viral genome chromatin assembly and/or maintenance occurs in the vDCP-NBs, some of our data tend to show that it unlikely represents the only pathway for chromatinization of the incoming naked HSV-1 genomes. Indeed, the depletion of DAXX, ATRX, UBN1 and to a lesser extent HIRA, significantly impacts the co-localization of the latent/quiescent HSV-1 genomes with PML, and hence the formation of vDCP-NBs, but only mildly affects the H3.3 association with the analyzed viral genome loci. However, beyond a compensatory mechanism between the two complexes that could bypass the requirement of the vDCP-NBs, we cannot exclude the possible contribution of other PML-NB-associated proteins involved in H3.3 chromatinization, as shown for the DEK protein [61]. Moreover, our data show that the depletion of DAXX, ATRX, HIRA, or UBN1 does not modify the accumulation of e-H3.3 at PML-NBs, leaving intact the upstream requirement of H3.3 accumulation in PML-NBs for H3.3-dependent viral chromatin assembly. We have recently shown that vDCP-NBs are dynamic structures that can fuse during the course of a latent infection [16]. It is thus possible that incoming viral genomes can be dynamically associated with vDCP-NBs to be chromatinized, and in the absence of one of the components of either H3.3 chaperone complex, this dynamic can be perturbed, resulting in some viral genomes that are not co-localized with PML. Given that depletion of none of the four proteins affects the structure of the PML-NBs, and considering the essential role of PML-NBs in the H3.3 chromatinization of the viral genomes, this possibility cannot be ruled out.

Altogether, our study demonstrates the essential role of a PML-NB/H3.3/H3.3 chaperone axis in the process of chromatinization of viral genomes adopting a vDCP-NB pattern, which represents an essential structural and functional aspect of HSV-1 latency establishment.

## Materials and Methods

### Ethics Statement

All procedures involving experimental animals conformed to the ethical standards of the Association for Research in Vision and Ophthalmology (ARVO) statement for the use of animals in research and were approved by the local Ethics Committee of the Institute for Integrative Biology of the Cell (I2BC) and the Ethics Committee for Animal Experimentation (CEEA) 59 (Paris I) under number 2012-0047 and in accordance with European Community Council Directive 2010/63/EU. For animal experiments performed in the USA: animals were housed in American Association for Laboratory Animal Care-approved housing with unlimited access to food and water. All procedures involving animals were approved by the Children’s Hospital Animal Care and Use Committee and were in compliance with the Guide for the Care and Use of Laboratory Animals (protocol number: IAUC2013-0162 of 2/28/2107 (yearly annual approval)).

### Virus strains, mice and virus inoculation: primary mouse TG neuron cultures, cells

The HSV-1 SC16 strain was used for mouse infections and has been characterized previously [62]. The HSV-1 mutant *in*1374 is derived from the 17 *syn* + strain and expresses a temperature-sensitive variant of the major viral transcriptional activator ICP4 [63] and is derived from *in*1312, a virus derived from the VP16 insertion mutant *in*1814 [64], which also carries a deletion/frameshift mutation in the ICP0 open reading frame [65] and contains an HCMV-*lacZ* reporter cassette inserted into the UL43 gene of *in*1312 [66]. This virus has been used and described previously [16,38]. All HSV-1 strains were grown in baby hamster kidney (BHK-21, American Type Culture Collection, ATCC CCL-10) cells and titrated in human osteosarcoma (U2OS, ATCC HTB-96) cells. *In*1374 was grown and titrated at 32°C in the presence of 3 mM hexamethylene bisacetamide [67]. PML wild-type, and knockout mice were obtained from the NCI Mouse Repository (NIH, <u>http://mouse.ncifcrf.gov</u>; strain, 129/Sv-*Pml^tm1Ppp^*) [68]. Genotypes were confirmed by PCR, according to the NCI Mouse Repository guidelines with primers described in [15].

Mice were inoculated and TG processed as described previously [15] Maroui et al., 2016}. Briefly, for the lip model: 6-week-old inbred female BALB/c mice (Janvier Labs, France) were inoculated with 10^6^ PFU of SC16 virus into the upper-left lip. Mice were sacrificed at 6 or 28 dpi. Frozen sections of mouse TG were prepared as described previously [15,69]. For the eye model: inoculation was performed as described previously [70]. Briefly, prior to inoculation, mice were anesthetized by intra-peritoneal injection of sodium pentobarbital (50 mg/kg of body weight). A 10-µL drop of inoculum containing 10^5^ PFU of 17syn+ was placed onto each scarified corneal surface. This procedure results in ~80% mouse survival and 100% infected TG.

Primary mouse TG neuron cultures were established from OF1 male mice (Janvier lab), following a previously described procedure {Maroui et al., 2016}. Briefly, 6–8-week-old mice were sacrificed before TG removal. TG were incubated at 37°C for 20 min in papain (25 mg) (Worthington) reconstituted with 5 mL Neurobasal A medium (Invitrogen) and for 20 min in Hank’s balanced salt solution (HBSS) containing dispase (4.67 mg/mL) and collagenase (4 mg/mL) (Sigma) on a rotator, and mechanically dissociated. The cell suspension was layered twice on a five-step OptiPrep (Sigma) gradient, followed by centrifugation for 20 min at 800 *g*. The lower ends of the centrifuged gradient were transferred to a new tube and washed twice with Neurobasal A medium supplemented with 2% B27 supplement (Invitrogen) and 1% penicillin–streptomycin (PS). Cells were counted and plated on poly-D-lysine (Sigma)- and laminin (Sigma)-coated, eight-well chamber slides (Millipore) at a density of 8,000 cells per well. Neuronal cultures were maintained in complete neuronal medium consisting of Neurobasal A medium supplemented with 2% B27 supplement, 1% PS, L-glutamine (500 µM), nerve growth factor (NGF; 50 ng/mL, Invitrogen), glial cell-derived neurotrophic factor (GDNF; 50 ng/mL, PeproTech), and the mitotic inhibitors fluorodeoxyuridine (40 µM, Sigma) and aphidicolin (16.6 µg/mL, Sigma) for the first 3 days. The medium was then replaced with fresh medium without fluorodeoxyuridine and aphidicolin.

Primary human foreskin (BJ, ATCC, CRL-2522), lung (IMR-90, Sigma, 85020204), fetal foreskin (HFFF-2, European Collection of Authenticated Cell Cultures, ECACC 86031405, kind gift from Roger Everett, CVR-University of Glasgow) fibroblast cells, primary human hepatocyte (HepaRG, HPR101, kind gift from Olivier Hantz & Isabelle Chemin, CRCL, Lyon, France) cells, human embryonic kidney (HEK 293T, ATCC CRL-3216) cells, U2OS, mouse embryonic fibroblast (MEF) *pml* +/+, MEF *pml* -/- cells (kind gift from Valérie Lallemand, Hopital St Louis, Paris), and BHK-21 cells were grown in Dulbecco’s Modified Eagle’s Medium (DMEM) supplemented with 10% fetal bovine serum (Sigma, F7524), L-glutamine (1% v/v), 10 U/mL penicillin, and 100 mg/mL streptomycin. BJ cell division is stopped by contact inhibition. Therefore, to limit their division, cells were seeded at confluence before being infected at a multiplicity of infection (m.o.i.) of 3, and then maintained in 2% serum throughout the experiment. Infections of BJ cells for short times (from 30 min to 6 h) were performed by synchronizing the infection process with a pre-step of virus attachment to the cells at 4°C for one hour. The infection medium was then removed, and the temperature was shifted to 37°C to allow a maximum of viruses to simultaneously penetrate into the cells.

### Frozen sections

Frozen sections of mouse TG were generated as previously described [69]. Mice were anesthetized at 6 or 28 d.p.i., and before tissue dissection, mice were perfused intra-cardially with a solution of 4% formaldehyde, 20% sucrose in 1X PBS. Individual TG were prepared as previously described [69], and 10-µm frontal sections were collected in three parallel series and stored at - 80°C.

### DNA-FISH and immuno-DNA FISH

HSV-1 DNA FISH probes consisting of cosmids 14, 28 and 56 [71] comprising a total of ~90 kb of the HSV-1 genome were labeled by nick-translation (Invitrogen) with dCTP-Cy3 (GE Healthcare) and stored in 100% formamide (Sigma). The DNA-FISH and immuno-DNA FISH procedures have been described previously [15,69]. Briefly, infected cells or frozen sections were thawed, rehydrated in 1x PBS and permeabilized in 0.5% Triton X-100. Heat-based unmasking was performed in 100 mM citrate buffer, and sections were post-fixed using a standard methanol/acetic acid procedure and dried for 10 min at RT. DNA denaturation of the section and probe was performed for 5 min at 80°C, and hybridization was carried out overnight at 37°C. Sections were washed 3 × 10 min in 2 x SSC and for 3 × 10 min in 0.2 x SSC at 37°C, and nuclei were stained with Hoechst 33258 (Invitrogen). All sections were mounted under coverslips using Vectashield mounting medium (Vector Laboratories) and stored at 4°C until observation.

For immuno-DNA FISH, cells or frozen sections were treated as described for DNA-FISH up to the antigen-unmasking step. Tissues were then incubated for 24 h with the primary antibody. After three washes, secondary antibody was applied for 1 h. Following immunostaining, the cells were post-fixed in 1% PFA, and DNA FISH was carried out from the methanol/acetic acid step onward. The same procedures were used for infected neuronal cultures except that the cells were fixed in 2% PFA before permeabilization.

### Western blotting

Cells were collected in lysis buffer (10 mM Tris-EDTA, pH 8.0) containing a protease inhibitor cocktail (Complete EDTA-free; Roche) and briefly sonicated. Protein extracts were homogenized using QiaShredders (Qiagen). The protein concentration was estimated by the Bradford method. Extracted proteins were analyzed by Western blotting using appropriate antibodies (see below).

### Microscopy, imaging, and quantification

Observations and most image collections were performed using an inverted Cell Observer microscope (Zeiss) with a Plan-Apochromat ×100 N.A. 1.4 objective and a CoolSnap HQ2 camera from Molecular Dynamics (Ropper Scientific) or a Zeiss LSM 800 confocal microscope. Raw images were processed using ImageJ software (NIH).

### Lentivirus and retrovirus production and establishment of cell lines

BJ or MEF cell lines expressing H3.1-SNAP-HAx3 (e-H3.1), H3.3-SNAP-HAx3 (e-H3.3), or Myc-hDAXX were established by retroviral transduction [72]. Briefly, pBABE plasmids encoding H3.1-SNAP-HAx3 or H3.3-SNAP-HAx3 (gift from Dr Jansen), pLNCX2 encoding Myc-hDAXX [36], were co-transfected with pCL-ampho (for subsequent transduction of BJ cells) or pCL-eco (for subsequent transduction of MEF cells) plasmids [73] by the calcium phosphate method into HEK 293T cells to package retroviral particles [74]. BJ cells stably expressing HIRA-HA and HA-UBN1 or transiently expressing the shRNAs were established by lentiviral transduction. Briefly, pLenti encoding HIRA-HA or HA-UBN1, pLKO empty, pLKO shPML_01, 02, shDAXX_01, 02, shATRX_01, 02, shHIRA_01, 02, shUBN1_01, 02, were co-transfected with psPAX.2 and pMD2.G plasmids by the calcium phosphate method into HEK 293T cells to package lentiviral particles. After 48 h, supernatant containing replication-incompetent retroviruses or lentiviruses was filtered and applied for 24 h on the target BJ or MEF cells in a medium containing polybrene 8 μg/mL (Sigma) [72]. Stable transfectants were selected with Blasticidin S (5 μg/mL, Sigma), puromycin (1 μg/mL, Sigma), or neomycin (G418, 1 mg/mL, Millipore) for 3 days, and a polyclonal population of cells was used for all experiments.

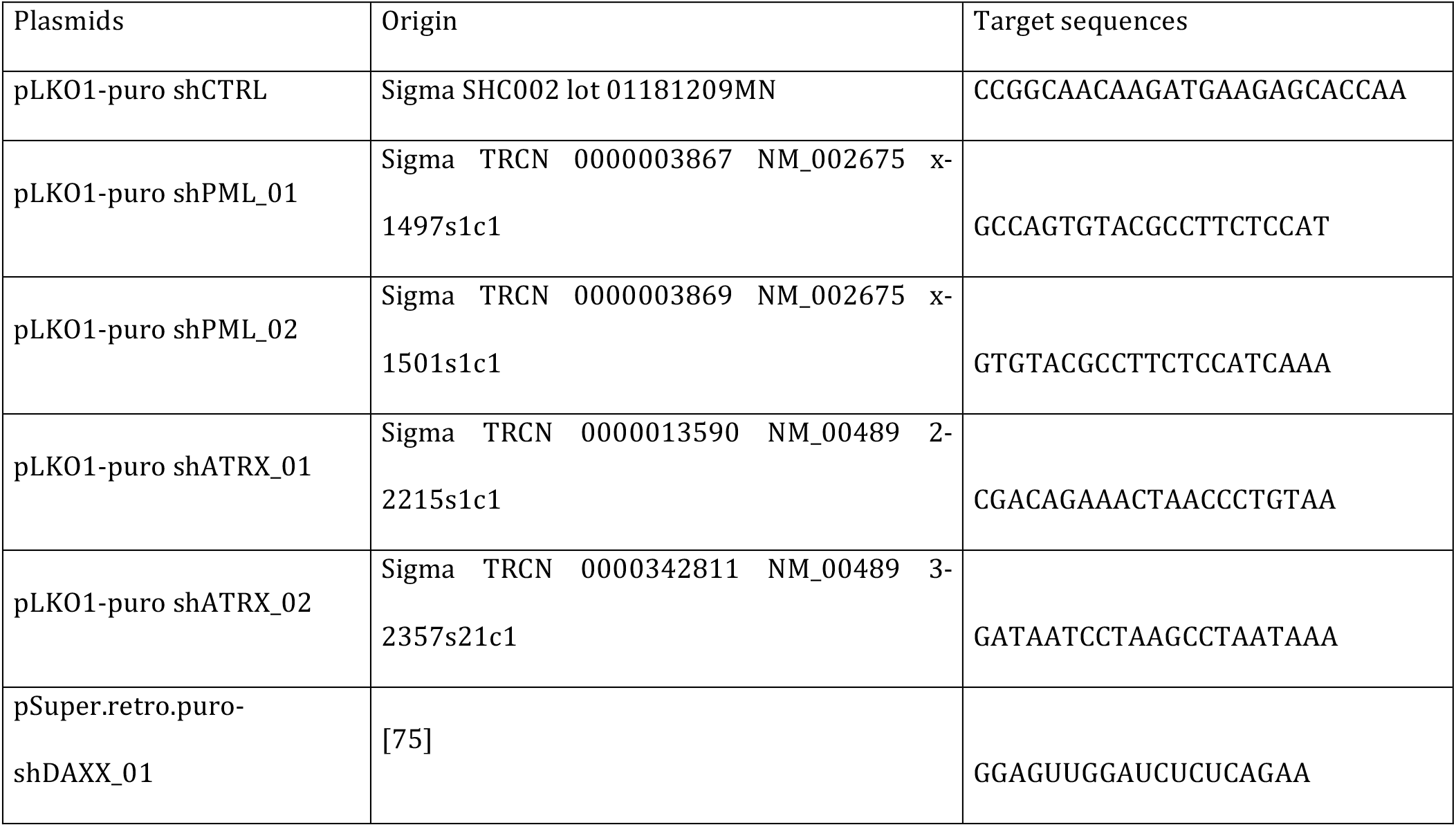
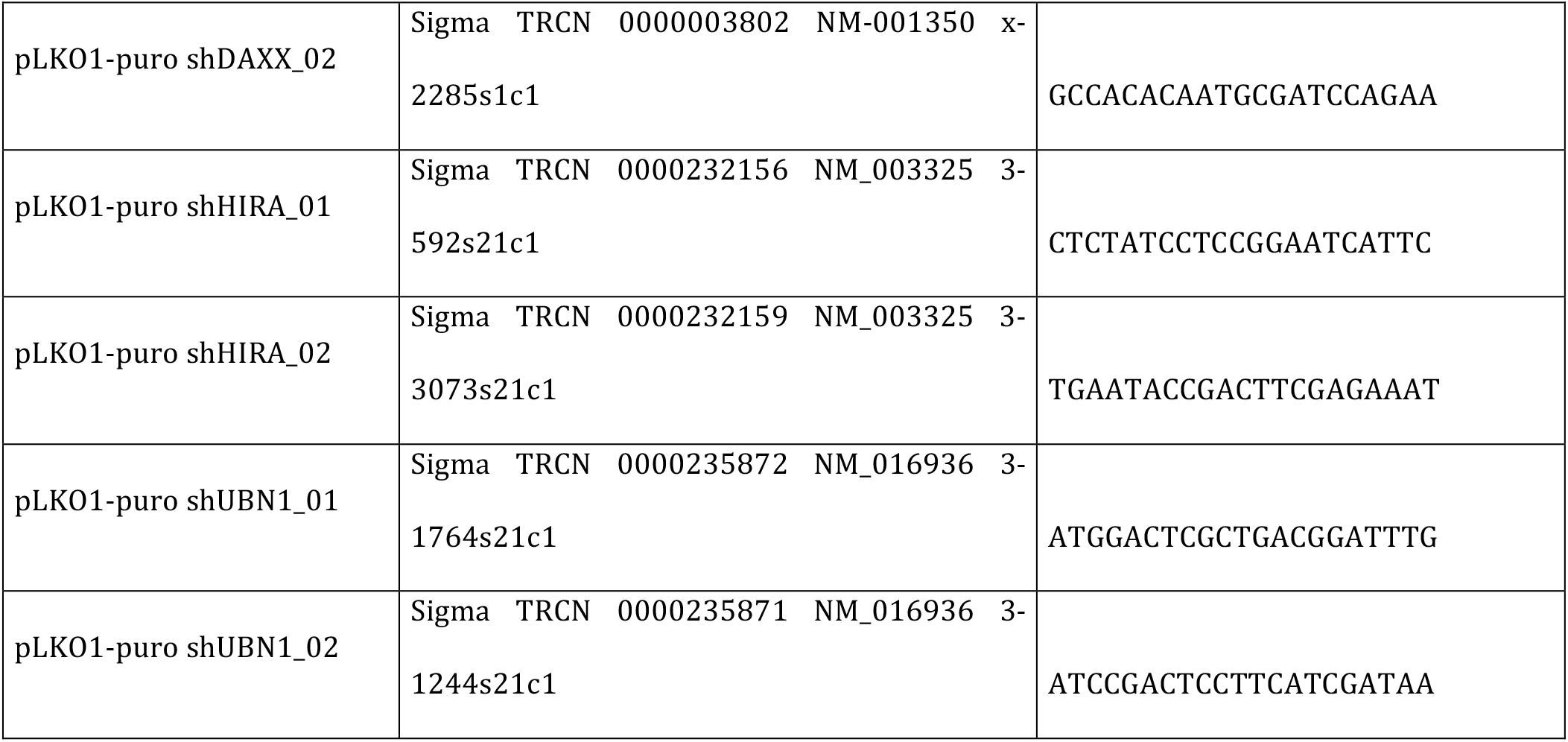

### ChIP and quantitative PCR

Cells were fixed with formaldehyde 2% for 5 min at RT, and then glycine 125 mM was added to arrest fixation for 5 min. After two washes with ice-cold PBS, the cells were resuspended in buffer A (100 mM Tris HCl pH 9.4, 10 mM DTT) and subjected to two 15 min-incubations on ice then at 30°C. The cells were subsequently lysed in lysis buffer B (10 mM Hepes pH 6.5, 0.25 % Triton, 10 mM EDTA, 0.5 mM EGTA) for 5 min at 4°C to recover the nuclei. Nuclei were incubated for 5 min at 4°C in buffer C (10 mM Hepes pH 6.5, 2 M NaCl, 10 mM EDTA, 0.5 mM EGTA) and then resuspended in 380 µl of buffer D (10 mM EDTA, 50 mM Tris HCl pH 8, 1% SDS, Protease Inhibitor Cocktail (Complete EDTA-free; Roche)). Nuclei were sonicated with a Bioruptor (Diagenode) for 30 cycles of 30 sec. “ON”, 30 sec. “OFF”. Thirty microliters of the sonication product was kept for the input, 50 µL for analyzing the sonication efficiency, and 300 µl diluted 10 times in RIPA buffer (50 mM Tris-HCl pH 8, 150 mM NaCl, 2 mM EDTA pH 8, 1% NP40, 0.5% Na Deoxycholate, 0.1% SDS, PIC 1x) for ChIP. Two micrograms of Ab was added and incubated overnight at 4°C. Sixty microliters of agarose beads coupled to protein A (Millipore 16-157) or G (Millipore 16-201) was added for 2 h at 4°C under constant shaking. Beads were then successively washed for 5 min at 4°C under constant shaking once in “low salt” (0.1% SDS, 1% Triton X-100, 2 mM EDTA, 20 mM Tris HCl pH 8.0, 150 mM NaCl) buffer, once in “high salt” (0.1% SDS, 1% Triton X-100, 2 mM EDTA, 20 mM Tris HCl pH 8.0, 500 mM NaCl) buffer, once in “LiCl” (0.25 mM LiCL, 1% NP40, 1% NaDOC, 1 mM EDTA, 10 mM Tris HCl pH 8.0) buffer and twice in TE (10 mM Tris pH 8.0, 1 mM EDTA) buffer. Chromatin-antibody complexes are then eluted twice at RT for 20 min under constant shaking with 250 µl of elution buffer (1% SDS, 0.1 M NaHCO3). Input and IP products were de-crosslinked overnight at 65°C and then treated for 2 h at the same temperature with 20 mg/mL of proteinase K (Sigma) and 10 mg/mL of RNAse A (Sigma). DNA was then purified by phenol-chloroform/ethanol precipitation, resuspended in water, and kept at -20°C until use for qPCR or ChIPseq.

Quantitative PCR was performed using Quantifast SYBR Green mix (Stratagene) and the MX3005P apparatus. Primers were used at a final concentration of 1 µM and were as follows:

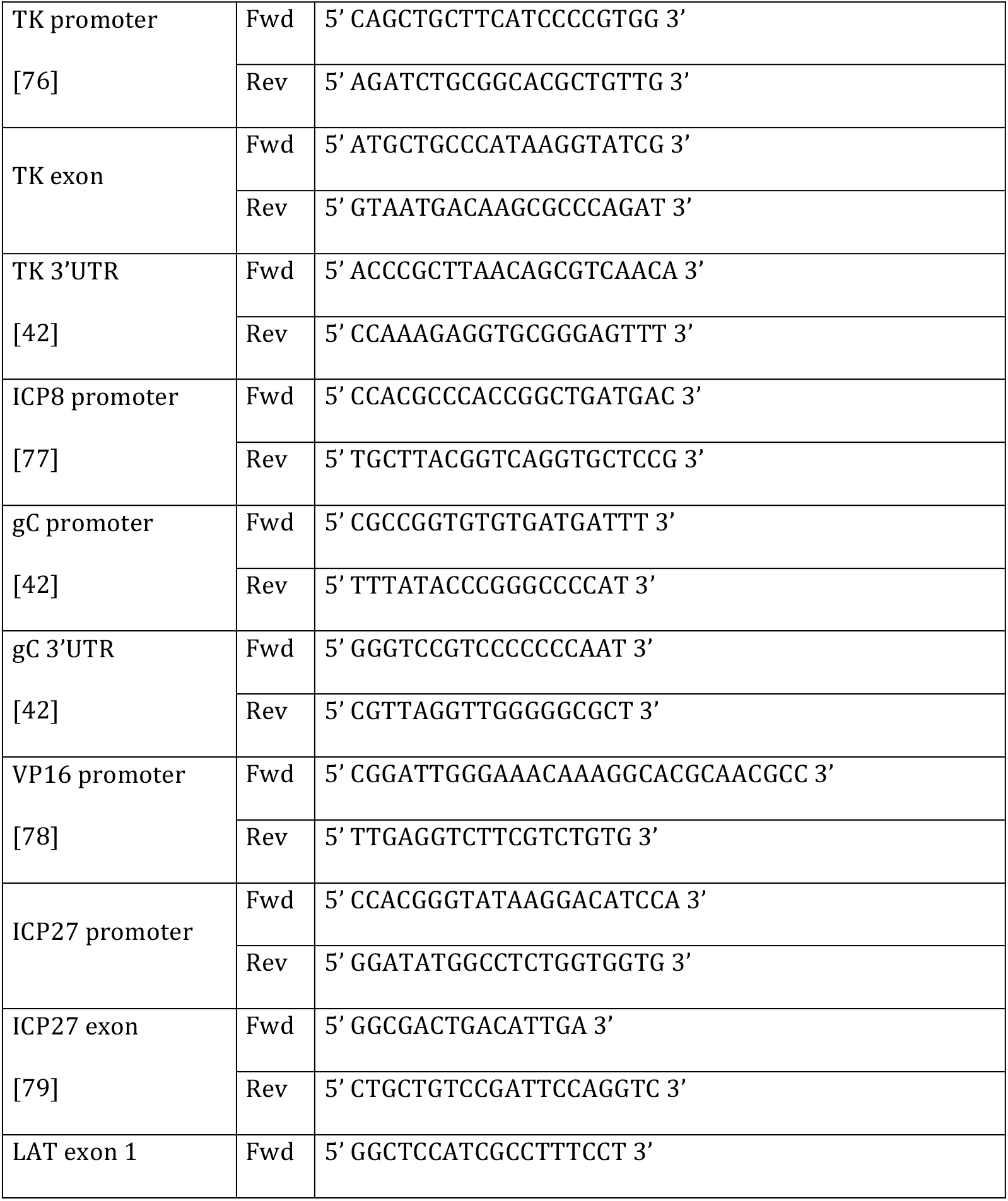
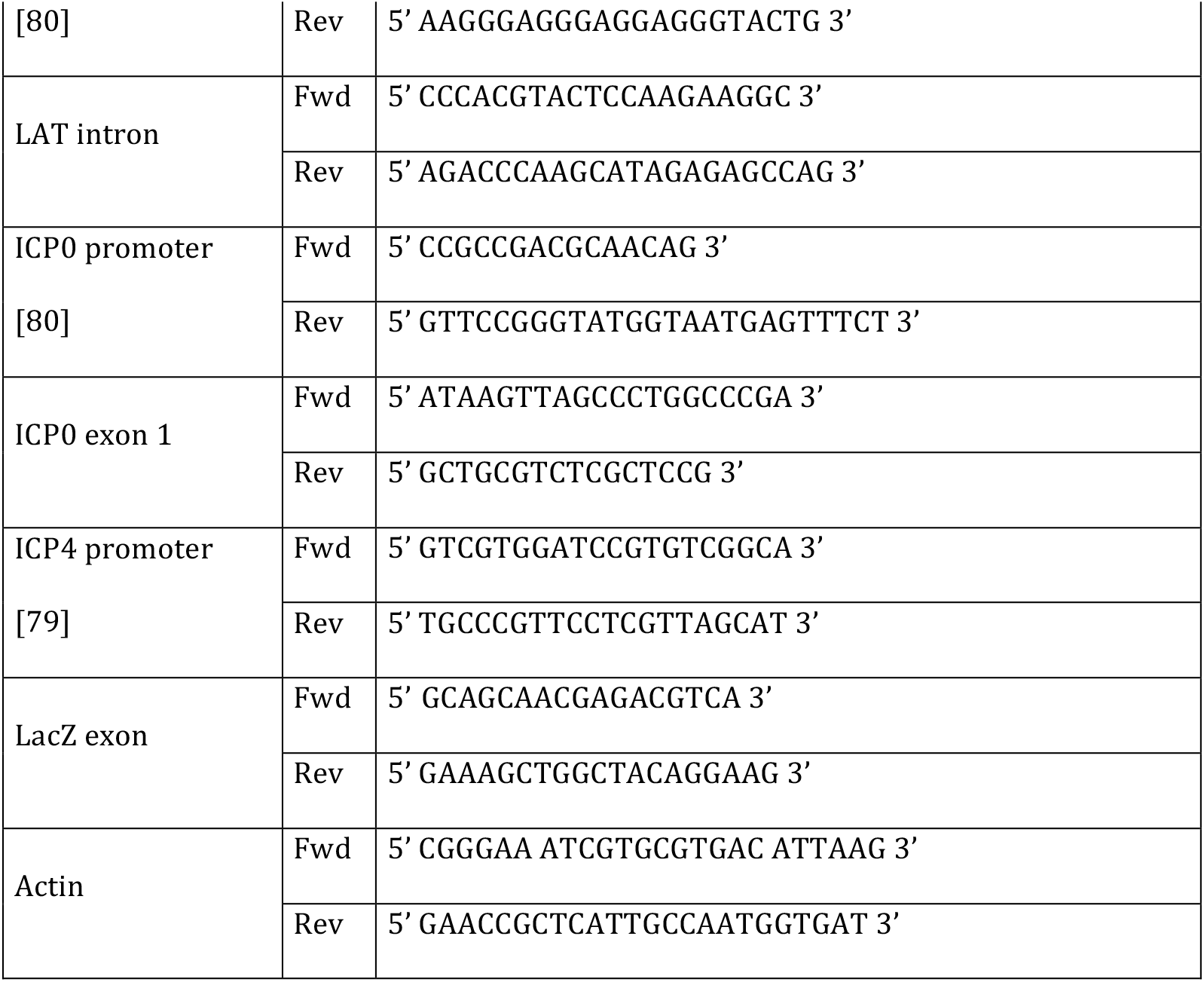

### siRNA transfections

Transfections of BJ cells with siRNAs was performed using Lipofectamine RNAiMAX and following the supplier’s procedure (Thermo Fisher Scientific). The following siRNAs were used at a final concentration of 40 nM for 48 h: siH3F3A: 5′-CUACAAAAGCCGCUCGCAA [81]; siH3F3B: 5′-GCUAAGAGAGUCACCAUCA [81].

### Antibodies

The following primary antibodies were used:

For immunofluorescence

*For immunofluorescence*
| Proteins | Origin | References | Species | Dilution |
| --- | --- | --- | --- | --- |
| Asf1a | Geneviève Almouzni<br>(Institut Curie, Paris) |  | Rabbit polyclonal | 1/1000 |
| ATRX (h-300) | Santa Cruz | sc-15408 | Rabbit polyclonal | 1/100 |
| c-Myc (9E10) | Santa cruz | sc-40 | Mouse monoclonal | 1/200 |
| Cabin1 | Sigma | HPA043296 | Rabbit polyclonal | 1/100 |
| DAXX (M-112) | Santa Cruz | sc-7152 | Rabbit polyclonal | 1/100 |
| HA | Abcam | AB9110 | Rabbit polyclonal | 1/500-1000 |
| HA (3F10) | Roche | 1867423 | Rat monoclonal | (1/1000) |
| HIRA | Abcam | ab20655 | Rabbit polyclonal | 1/100 |
| HIRA (WC119) | Millipore | 04-1488 | Mouse monoclonal | 1/100 |
| p48 (CAF-1) | Abcam | ab1765 | Rabbit polyclonal | 1/100 |
| p60 (CAF-1) | Novus | 500-207 | Mouse monoclonal | 1/100 |
| p150 (CAF-1) | Novus | 500-212 | Mouse monoclonal | 1/100 |
| human PML (H-238) | Santa Cruz | sc-5621 | Rabbit polyclonal | 1/500 |
| human PML(PG-M3) | Santa Cruz | sc-966 | Mouse monoclonal | 1/500 |
| human PML 5E10 | Roel Van Driel<br>(University of<br>Amsterdam) |  | Mouse monoclonal | 1/500 |
| mouse PML | Millipore | MAB3738 | Mouse monoclonal | 1/100 |
| Sp100 | Thomas Sternsdorf<br>(University of Hamburg) | SpGH | Rabbit polyclonal | 1/500 |
| SUMO-1 | Cell Signaling | 4930 | Rabbit polyclonal | 1/500 |
| SUMO-1 (5B12) | MBL | M113-3 | Mouse monoclonal | 1/500 |
| SUMO-2/3 (18H8) | Cell Signaling | 4971 | Rabbit monoclonal | 1/500 |
| SUMO-2/3 (1E7) | MBL | M114-3 | Mouse monoclonal | 1/500 |
| UBN1 Zap1 | Henri Gruffat<br>(CIRI, ENS, Lyon) |  | Mouse monoclonal | 1/100 |

All secondary antibodies were Alexa Fluor-conjugated and were raised in goats (Invitrogen).

For ChIP

All secondary antibodies were Alexa Fluor-conjugated and were raised in goats (Invitrogen). *For ChIP*
| Proteins | Origin | References | Species |
| --- | --- | --- | --- |
| ATRX (h-300) | Santa Cruz | sc-15408 | Rabbit polyclonal |
| c-Myc (9E10) | Millipore | MABE282 | Mouse monoclonal |
| H3.1/2 | Millipore | ABE154 | Rabbit polyclonal |
| H3.3 | Millipore | 09-838 | Rabbit polyclonal |
| HA | ABCAM | AB9110 | Rabbit polyclonal |
| IgG Rabbit | Diagenode | Kch-504-250 | Rabbit polyclonal |

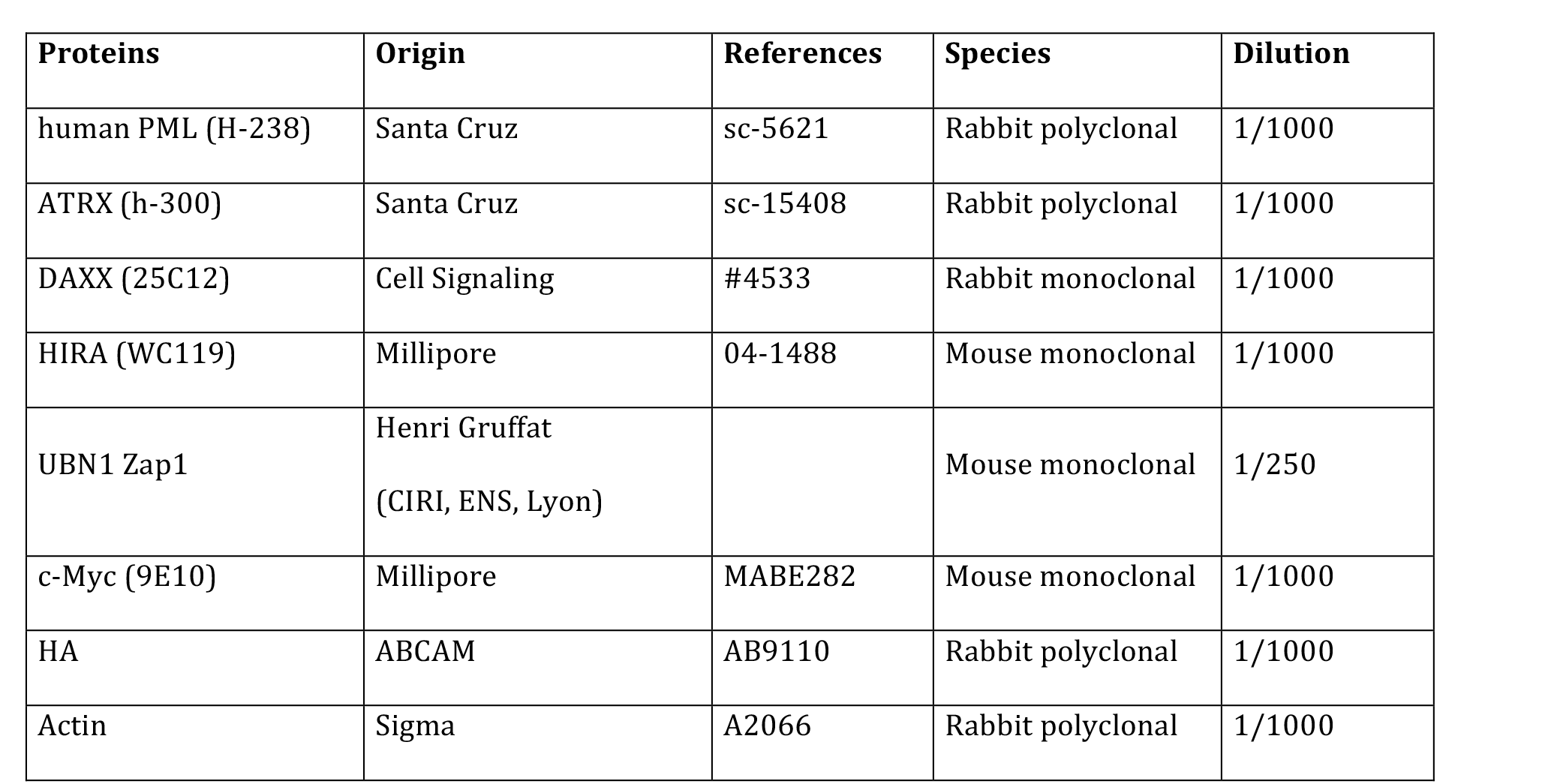
For WB

All secondary antibodies were HRP-conjugated and were raised in goats (Sigma).

## Acknowledgements

We thank Roel van Driel (University of Amsterdam, Netherland) for the PML antibody (mAb 5E10), Henri Gruffat (CIRI, ENS Lyon, France) for the UBN1 antibody, Geneviève Almouzni (Institut Curie, Paris, France) for the ASF1a antibody, Thomas Sternsdorf (University of Hamburg, Germany) for the Sp100 antibody, Roger Everett and Chris Preston (Center for Virus Research, University of Glasgow, UK) for the *in*1374 virus, Wade Bresnahan (University of Texas, USA) for the pSuper.retro.puro-shDAXX_01, Lars Jansen (Insituto Gulbenkian de Ciencia, Portugal) for the pBABE plasmids encoding H3.1-SNAP-HAx3 and H3.3-SNAP-HAx3, Olivier Hantz and Isabelle Chemin (CRCL, Lyon) for the HepaRG cells, Valérie Lallemand-Breitenbach (Hopital Saint Louis, Paris, France) for the MEF *pml*^+/+^ and *pml*^−/-^ cells and the Centre Technologique des Microstructures (CTµ) of the Université Claude Bernard Lyon 1 for the confocal microscopy.

## Supplementary Figures

Figure S1. Latent/quiescent HSV-1 genomes co-localize with PML and PML-NB-associated proteins in vDCP-NBs.

Data from immuno-FISH experiments performed in human primary fibroblasts (BJ cells) infected for 2 days with the replication-defective HSV-1 virus *in*1374. PML (i), Sp100 (ii), SUMO-1 (iii), SUMO 2/3 (iv), ATRX (v), DAXX (vi) (green), and HSV-1 genomes (red) were detected. Scale bars = 5 µm.

Figure S2. Expression of tagged versions of DAXX (Myc), HIRA and UBN1 (HA) in BJ cells.

Normal BJ cells were transduced with lentiviruses expressing Myc-DAXX, HIRA-HA, or HA-UBN1, and stable cell lines expressing the tagged proteins were selected by puromycin selection. Expression of the tagged proteins was detected by immunofluorescence (A) and Western blotting (B). For WB, actin was used as a loading control. Scale bars = 5 µm.

Figure S3. Expression of the tagged H3.3 (e-H3.3) and H3.1 (e-H3.1) in BJ cells.

(A) Detection of the protein expression by WB using the anti-HA antibody. Endogenous histone H3 was detected as a control.

Actin was detected as a loading control.

(B) Detection of e-H3.1 (i) and e-H3.3 (ii) by immunofluorescence.

(C) Co-detection of e-H3.1 (i) and e-H3.3 (ii) (green) with PML (red). E-H3.3, unlike e-H3.1, co-localizes with PML-NBs. Scale bars = 5 µm.

(D) Quantification of the immunofluorescence experiments performed in (C). Data from two independent experiments.

Figure S4. The native histone variant H3.3, but not the canonical H3.1/2, associates with latent/quiescent HSV-1 genomes.

(A) ChIP-qPCR performed in *in*1374-infected normal BJ cells using control IgG (blue), anti-H3.1/2 (red), or anti-H3.3 (green) antibodies.

(B) ChIP-qPCR performed in *in*1374-infected BJ e-H3.3 cells using control IgG (blue), anti-H3.1/2 (red), anti-H3.3 (green), or anti-HA (purple) antibodies.

Figure S5. Validation of the shRNAs against DAXX, ATRX, HIRA, and UBN1.

(A) BJ cells were transduced with shRNA-expressing lentiviruses before analysis 48 h post-transduction. RT-qPCR to detect DAXX, ATRX, HIRA, and UBN1 mRNA was performed, and the results were compared to a control shRNA (shCTRL). Data represent means from three independent experiments ± SD. The Student’s *t*-test was applied to assess the significance of the results. * = p< 0.05, ** = p< 0.01.

(B) WB for detection of decreases in DAXX, ATRX, HIRA, and UBN1 proteins (48 h post-transduction) in normal BJ cells or BJ cells transduced with shRNA-expressing lentiviruses. Actin was detected as a loading control. Two shRNAs were tested for each protein.

Figure S6. Effects of the depletion of DAXX, ATRX, HIRA or UBN1 on PML-NB detection. BJ cells were transduced with shRNA-expressing lentiviruses before analysis 48 h post-transduction. Immunofluorescences were performed to detect DAXX, ATRX, HIRA, and UBN1 (green) and PML (gray, red). Two shRNAs were tested for each protein. Scale bars = 5 µm.

Figure S7. Validation of the shRNAs against DAXX, ATRX, HIRA, and UBN1 in e-H3.3-expressing BJ cells. H3.3-expressing BJ cells were transduced with shRNA-expressing lentiviruses before analysis 48 h post-transduction.

(A) RT-qPCR to quantify DAXX, ATRX, HIRA, and UBN1 mRNA was performed, and the results were compared to a control shRNA (shCTRL). Data represent means from two independent experiments.

(B) WB for detection of decreases in ATRX, HIRA, and UBN1 proteins (48 h post-transduction) in normal e-H3.3-expressing BJ cells or e-H3.3-expressing BJ cells transduced with shRNA-expressing lentiviruses. Actin was detected as a loading control.

Figure S8. Depletion of DAXX, ATRX, HIRA, or UBN1 does not affect the accumulation of e-H3.3 in PML-NBs. Immunofluorescence experiments performed in e-H3.3-expressing BJ cells transduced with a lentivirus expressing a control shRNA (shCTRL, i, iii, v, vii) or a shRNA targeting DAXX (ii), ATRX (iv), HIRA (vi), or UBN1 (viii). E-H3.3 (gray, green); DAXX, ATRX, HIRA, UBN1 (gray, red); and PML (gray, blue) were detected. For the staining, a rat anti-HA mAb was used to detect e-H3.3, a rabbit polyclonal for the detection of DAXX, ATRX (i-iv), or PML (v-viii), and a mouse mAb for the detection of HIRA, UBN1 (v-viii) or PML (i-iv). Arrowheads note an example of e-H3.3 co-localization with PML-NBs in each sample. Scale bars = 5 µm.

Figure S9. Validation of the shRNAs against PML in normal and e-H3.3-expressing BJ cells.

Normal (A-C) or e-H3.3-expressing (D and E) BJ cells were transduced with a lentivirus expressing a control shRNA (shCTRL) or PML shRNAs (shPML) before analysis. Two different shRNAs were validated in normal BJ cells.

(A) Immunofluorescence to detect the PML-NB signal.

(B) RT-qPCR to quantify PML mRNA. Data represent means from three independent experiments ± SD. The Student’s *t*-test was applied to assess the significance of the results. * = p< 0.05, ** = p< 0.01.

(C) WB to detect PML protein.

(D) RT-qPCR to quantify PML mRNA. Data represent means from two independent experiments.

(E) WB to detect PML protein.

Figure S10. The decrease in the H3.3 association with latent/quiescent HSV-1 genomes in BJ cells depleted for PML-NBs is not compensated by H3.1. ChIP-qPCR for the detection of e-H3.1 associated with HSV-1 in e-H3.1-expressing BJ cells infected with *in*1374 for 24 h and previously transduced with a lentivirus expressing a shRNA control (shCTRL, blue) or a PML shRNA (shPML, red). Anti-HA antibody was used for the ChIP experiments. The analyzed viral loci were described previously. The data for ICP27pro and ICP0pro loci are not shown because the CTs in the PML shRNA samples were above the quantification range. Data represent means from two independent experiments ± SD.

Figure S11. The specific depletion of H3.3 does not affect the PML-NBs.

(A) WB to visualize the depletion of H3.3 in e-H3.3-expressing BJ cells. A combination of two siRNAs targeting H3.3 transcripts from both H3.3-encoding genes (H3F3A and H3F3B) were used for the depletion of H3.3. Actin was detected as a loading control.

(B) Immunofluorescence experiment performed in e-H3.3-expressing BJ cells transfected with control (siCTRL) or H3.3 (siH3F3A+3B) siRNAs. E-H3.3 (green), and PML (red) were detected. Nuclei were detected with DAPI (gray). Scale bars = 5 µm.

(C) Quantifications of co-localizations of HSV-1 genomes with PML issued from immuno-FISH experiments performed in in1374-infected BJ cells (2 dpi) previously transfected with control (siCTRL) or H3.3 (siH3F3A+3B) siRNAs. Data represent means from three independent experiments ± SD. The data suggest that vDCP-NBs are independent of H3.3 chromatinization of the latent/quiescent HSV-1 genomes for their formation.

